# A bacteriophage-based, highly efficacious, needle and adjuvant-free, mucosal COVID-19 vaccine

**DOI:** 10.1101/2022.04.28.489809

**Authors:** Jingen Zhu, Swati Jain, Jian Sha, Himanshu Batra, Neeti Ananthaswamy, Paul B. Kilgore, Emily K. Hendrix, Yashoda M. Hosakote, Xiaorong Wu, Juan P. Olano, Adeyemi Kayode, Cristi L. Galindo, Simran Banga, Aleksandra Drelich, Vivian Tat, Chien-Te K. Tseng, Ashok K. Chopra, Venigalla B. Rao

**Author notes:** shared first authors (Jingen Zhu, Swati Jain, Jian Sha). shared second authors (Himanshu Batra, Neeti Ananthaswamy).

## Abstract

The authorized mRNA- and adenovirus-based SARS-CoV-2 vaccines are intramuscularly injected and effective in preventing COVID-19, but do not induce efficient mucosal immunity, or prevent viral transmission. We developed a bacteriophage T4-based, multicomponent, needle and adjuvant-free, mucosal vaccine by engineering spike trimers on capsid exterior and nucleocapsid protein in the interior. Intranasal administration of T4-COVID vaccine induced higher virus neutralization antibody titers against multiple variants, balanced Th1/Th2 antibody and cytokine responses, stronger CD4^+^ and CD8^+^ T cell immunity, and higher secretory IgA titers in sera and bronchoalveolar lavage with no effect on the gut microbiota, compared to vaccination of mice intramuscularly. The vaccine is stable at ambient temperature, induces apparent sterilizing immunity, and provides complete protection against original SARS-CoV-2 strain and its Delta variant with minimal lung histopathology. This mucosal vaccine is an excellent candidate for boosting immunity of immunized and/or as a second-generation vaccine for the unimmunized population.

## INTRODUCTION

The mRNA, adenovirus-based, and inactivated viral vaccines currently used for human immunization are having a tremendous impact on tamping down the devastating COVID-19 pandemic that has caused millions of deaths across the globe. Administered by intramuscular injections, these vaccines remain as the major source for the rest of the world’s unvaccinated population. Many other vaccines are at various stages of preclinical studies and clinical trials (Tregoning et al., 2021). However, there are yet no needle-free mucosal vaccines authorized for human administration (Alu et al., 2022; Chavda et al., 2021).

Although the injectable vaccines are highly effective (70-95%) in preventing severe symptoms of the disease, hospitalization of patients, and deaths, they do not efficiently prevent viral acquisition or viral shedding from infected individuals. This is attributed to the lack of vaccine-induced secretory IgA (sIgA) mucosal immune responses in the respiratory airways that could prevent person-to-person transmission (Alu et al., 2022; Corbett et al., 2020; Mercado et al., 2020). Therefore, risk of transmission from vaccinated subjects, who are susceptible to SARS-CoV-2 infection, as seen currently on a global scale with the highly transmissible Omicron variants, remains a serious concern (Tiboni et al., 2021).

The current vaccines developed using the spike protein of the ancestral SARS-CoV-2 strain (Wuhan-Hu-1) show progressively diminished efficacy against the subsequently emerged viral variants of concern (VOCs) such as Alpha, Beta, Gamma, Delta, and most recently Omicron and its subvariant BA.2, which are more efficiently transmitted and/or more lethal. The evolutionary space for emergence of newer SARS-CoV-2 variants/subvariants that are even more efficiently transmissible and also more lethal that might render the current vaccines ineffective remains a worrisome and real possibility (Markov et al., 2022).

Considering the evolutionary path of the virus, the most desired next-generation vaccine(s) would be one that can induce strong mucosal immunity, in addition to broader systemic immunity (Alu et al., 2022; Chavda et al., 2021; Focosi et al., 2022; Lavelle and Ward, 2021). Elicitation of target-specific mucosal antibodies at the portal of virus entry would block virus acquisition as well as shedding of infectious virus particles and their potential transmission (Afkhami et al., 2022; Bricker et al., 2021; Hassan et al., 2020; Hassan et al., 2021; Ku et al., 2021; Sterlin et al., 2021b; van Doremalen et al., 2021b). Such platforms are of particular strategic importance at this stage of the COVID-19 pandemic. Additionally, platforms that are needle- and adjuvant-free and stable at ambient temperatures would greatly accelerate global distribution efforts, not only for controlling the current COVID-19 pandemic but also for any future epidemic or pandemic. Furthermore, needle-free vaccines can be administered easily and safely, and may provide the best option to vaccinate children.

We recently reported (Zhu et al., 2021) the development of a “universal” phage T4 vaccine design platform (Figures 1A) by <u>C</u>lustered <u>R</u>egularly Interspaced <u>S</u>hort <u>P</u>alindromic <u>R</u>epeats (CRISPR) engineering (Liu et al., 2020; Tao et al., 2017) that can rapidly generate multivalent vaccine candidates. Using an intramuscular immunization scheme, an optimal COVID-19 vaccine candidate (referred to as T4-CoV-2) was selected that elicited robust immunogenicity, virus neutralizing activity, and complete protection against ancestral SARS-CoV-2 challenge in a mouse model. This vaccine consisted of T4 phage decorated with ~100 copies of prefusion-stabilized spike ectodomain trimers (S-trimers) on the surface of 120 × 86 nm virus capsid (Fig. 1A). In addition, the vaccine also contained SARS-CoV-2 nucleocapsid protein (NP) packaged in the capsid core and a 12-amino acid (aa) peptide of the putative external domain of E protein (Ee) fused to the highly antigenic outer capsid protein (Hoc) displayed on the capsid surface (Fig. 1A).

**Figure 1.**
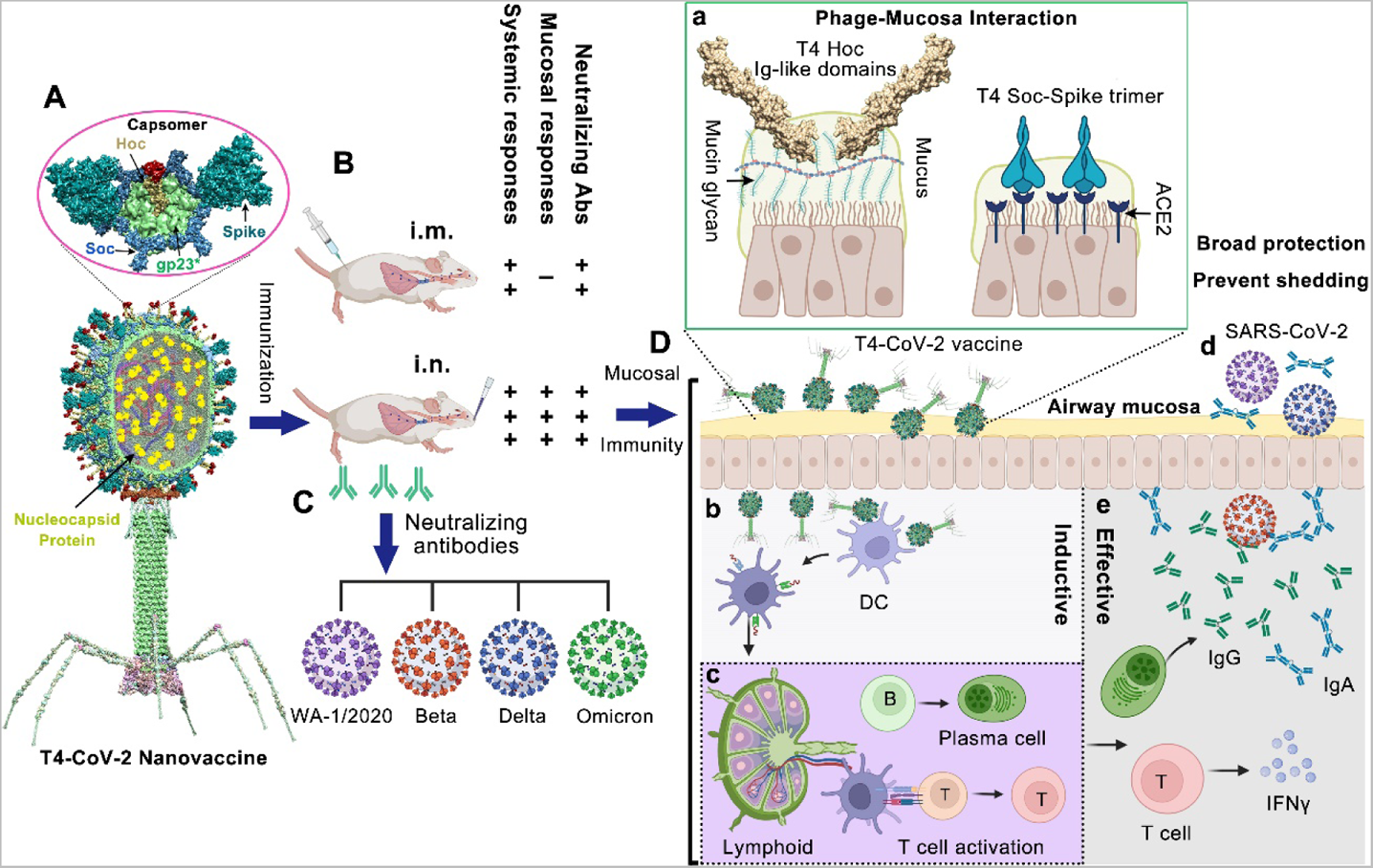
Intranasal vaccination of mice using bacteriophage T4-CoV-2 vaccine and possible mechanisms of protection. **(A)** Structural model of T4-CoV-2 nanovaccine constructed by CRISPR engineering (Zhu et al., 2021). The enlarged view shows a single hexameric capsomer consisting of six subunits of major capsid protein gp23* (green), trimers of Soc (blue), and a Hoc fiber (yellow) at the center of capsomer. The NP, Ee, and SpyCatcher gene were “hard-wired” by inserting the respective expressible genes into phage genome, which resulted in display of Ee peptide (red, 155 copies per T4) at the tip of Hoc fiber, SpyCatcher as Soc fusion on capsid surface (~200 copies per capsid), and packaging of NP molecules (yellow, 100 copies per T4) inside the capsid. The Spytagged Spike trimer (cyan) purified from ExpiCHO cells was then conjugated to Soc-SpyCatcher (Keeble et al., 2019). **(B and C)** Comparison between i.m. and i.n. T4-CoV-2 vaccination (B) and the elicited neutralizing antibodies against SARS-CoV-2 and its VOCs including Beta, Delta, and Omicron (C). **(D)** The mucosal immune responses induced by T4-CoV-2 i.n. vaccination. After i.n. inoculation, T4-CoV-2 particles would bind to mucosal cells: i) through the Ig-like domains of Hoc fibers which interact with mucin glycoproteins, and ii) through the displayed S-trimers which bind to ACE2 that is abundant in nasal epithelium **(a)**. Then, the antigen-presenting cells in the respiratory tract, such as dendritic cells (DCs) capture T4-CoV-2 phage **(b)**, migrate to mucosal-associated lymphoid tissues, and present the antigens to lymphocytes, including B and T cells **(c)**. The activated B cells become plasma cells secreting anti-SARS-CoV-2 IgG and IgA which neutralize virus within the respiratory tract **(d and e)**. The activated T cells migrate to lungs, produce cytokines and regulate the immune responses, and/or directly attack virus-infected host cells. These mucosal immune responses produced by T4-CoV-2 i.n. vaccination might be able to block viral entry (host’s viral acquisition) and viral exit (host’s viral shedding) in the respiratory tract.

The protective immunity of the T4-CoV-2 nanovaccine could potentially be because of the repetitive and symmetrical arrays of S-trimers on phage particles, resembling the PAMPs (pathogen-associated molecular patterns) present on human viral pathogens (de Vries et al., 2021; Freeman et al., 2021; Joyce et al., 2021; Tao et al., 2018b). This architecture might mimic, in some respects, the spikes displayed on the SARS-CoV-2 virion (Yao et al., 2020). Therefore, we hypothesized that it is probable that such a T4-CoV-2 nanoparticle when exposed to nasal mucosal surfaces might be recognized as a natural viral intruder by the resident immune cells, stimulating strong mucosal as well as systemic immune responses (Figures 1B to 1D). Furthermore, the S-trimer-displayed T4-CoV-2 nanoparticle could efficiently bind to the nasal epithelium that has the highest concentration of angiotensin-converting enzyme (ACE2) receptors (Hou et al., 2020). Additionally, the 155 symmetrically arranged Ig-like Hoc fibers on the T4 capsid are reported to interact with mucin glycoproteins, potentially capturing the T4-CoV-2 vaccine particles at the nasal mucosa (Barr Jeremy et al., 2015; Barr et al., 2013) (Figure 1D, a), translocation across the epithelial layer (Nguyen et al., 2017), and uptake by antigen-presenting cells (Popescu et al., 2021).

Here, we tested this hypothesis in a mouse model by intranasal (i.n.) inoculation of the T4-CoV-2 vaccine and compared the immune responses with those elicited by intramuscular (i. m.) injection. Remarkably, this needle- and adjuvant-free vaccination with non-infectious T4-COVID nanoparticles induced strong mucosal, humoral, and cellular immunity. The responses included spike-specific CD4^+^ helper and effector T cells and CD8^+^ killer T cells, and broad neutralization of SARS-CoV-2 VOCs including B.1.135 Beta, B.1.617.2 Delta, and B.1.1.529 Omicron, in both BALB/c as well as human ACE2 (hACE2) transgenic mouse models. Importantly, these responses elicited by needle-free vaccination are much stronger when compared to the injected vaccine, and strong mucosal secretory IgA antibodies were measured only in i.n.-vaccinated mice. Furthermore, the T4-CoV-2 vaccine is stable at ambient temperature, which can be easily manufactured and distributed at a modest cost. This phage-based mucosal vaccine, thus, is an excellent candidate for boosting the immunity of immunized and/or as a second-generation vaccine for the unimmunized populations.

## RESULTS

### Needle-free T4-CoV-2 nanovaccine stimulates more robust humoral and cellular immune responses against SARS-CoV-2 and VOCs than an injectable vaccine

The immunogenicity of the T4-CoV-2 nanovaccine was first evaluated in 5-week-old conventional BALB/c mice. In a standard prime-boost regimen (Figures 2A and 2B), animals received two i.m. or i.n. doses of either the T4 phage (vector control) or the T4-CoV-2 phage vaccine decorated with 20 µg (high-dose; ~2.5 × 10^11^ particles), 4.8 µg (medium-dose, ~6 × 10^10^ particles), or 0.8 µg (low-dose; ~1 × 10^10^ particles) of SARS-CoV-2 Spike-ectodomain (Secto, aa 1 - 1213) trimers. In a 1-dose regimen, animals received a single i.m. high-dose of the T4-CoV-2 vaccine.

**Figure 2.**
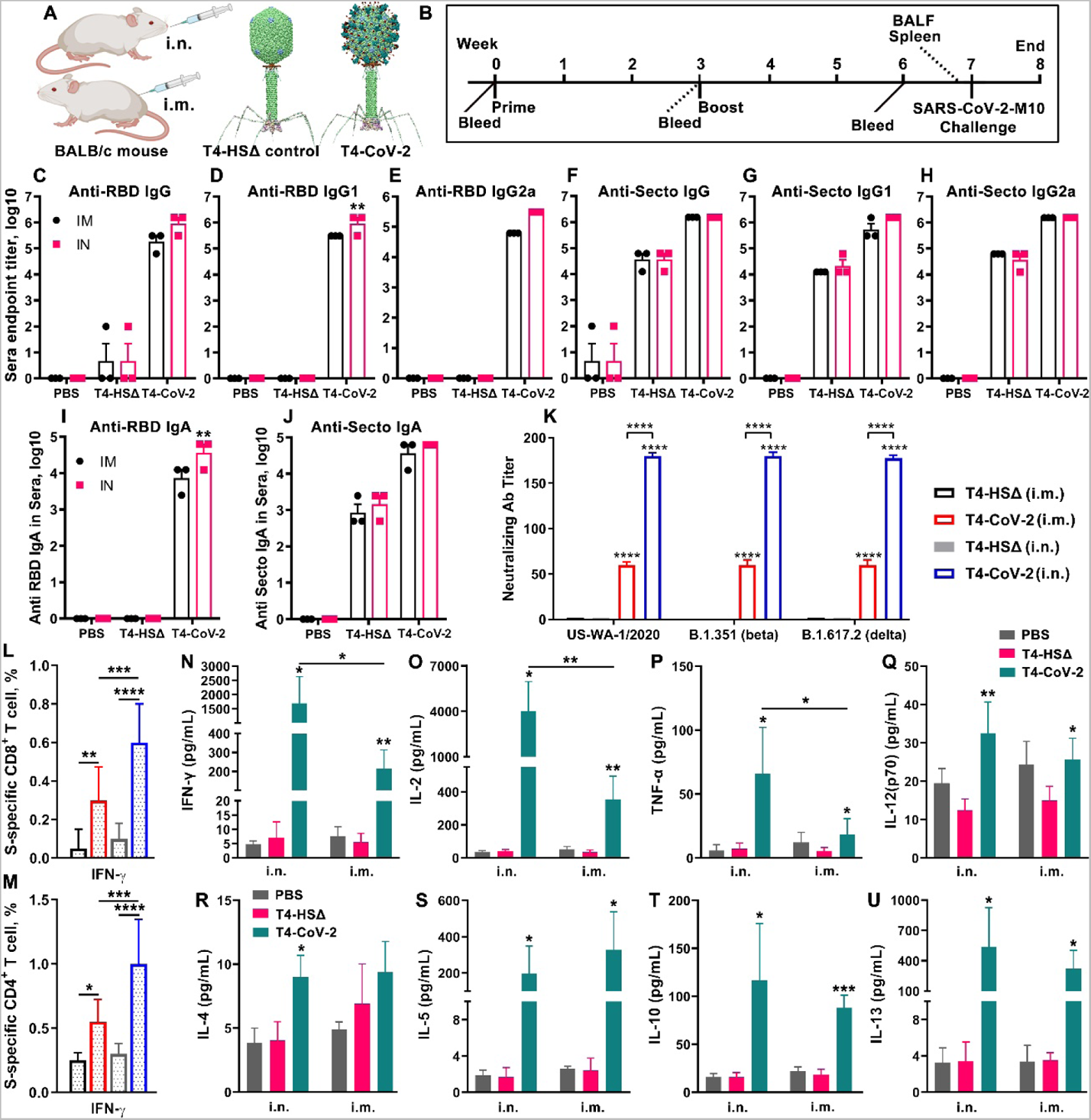
Intranasal immunization elicited greater anti-spike/RBD systemic humoral and cellular responses over intramuscular immunization. **(A)** Schematic of T4-CoV-2 i.n. and i.m. vaccinations with T4-HocΔ-SocΔ (T4-HSΔ) phage (left, vector control) and T4-CoV-2 recombinant phage (right, vaccine phage). **(B)** Scheme for vaccination and challenge. **(C to J)** Antibody responses in sera of immunized mice at day 21 after the last dose. Enzyme-linked immunosorbent assay (ELISA) was used to measure reciprocal endpoint antibody titers of anti-RBD IgG (C), anti-RBD IgG1 (D), anti-RBD IgG2a (E), anti-Secto IgG (F), anti-Secto IgG1 (G), anti-Secto IgG2a (H), anti-RBD IgA (I), and anti-Secto IgA (J). Data represent mean ± SEM. Data are from 3 pooled independent experiments (n = 22 for T4-CoV-2, n = 10 for T4-HSΔ, and n = 5 for PBS). **(K)** Virus neutralizing activity in sera of i.m. and i.n. immunized mice was determined by Vero E6 cell cytopathic assay using ancestral SARS-CoV-2 US-WA-1/2020, B.1.351 (Beta), and B.1.617.2 (Delta) strains. **(L and M)** Cellular immune responses. Percentages of IFNγ^+^ CD8^+^ (L) and IFNγ^+^ CD4^+^ (M) cells were plotted. **(N to U)** Cytokine responses. Representative Th1 (N to Q) and Th2 (R to U) cytokines are shown. For K to M, two-way ANOVA with Tukey post hoc test to compare multiple groups. For N to U, nonparametric Student’s t test to compare T4-vector control vs T4-CoV-2 vaccine groups and i.n. vs i.m. routes of vaccination. *P < 0.05; **P<0.01, ***P<0.001, ****P<0.0001. Data represent mean ± standard deviation and are representative of five biological replicates.

#### Antibody responses (IgG, isotypes, and IgA)

To evaluate humoral antibody responses, sera were collected on day 21 after the last dose (Figure 2B), and IgG, IgG1, and IgG2a antibodies specific to Secto protein or the receptor-binding domain (RBD) were quantified by ELISA (Figures 2C to 2H, Figure S1). The phosphate-buffered saline (PBS) and T4-vetor control groups, as expected, induced no significant antigen-specific antibodies, whereas the T4-CoV-2 vaccinated groups (either i.m. or i.n.) triggered high levels of IgG antibodies (Figures 2C and 2F).

High levels of both Th1 (IgG2a) and Th2 (IgG1) subtype antibodies were induced by i.m. and i.n. immunizations, demonstrating that the T4-CoV-2 vaccine triggered balanced Th1- and Th2-derived antibody responses (Figures 2D, 2E, 2G, and 2H). This is in contrast to the alum-adjuvanted subunit vaccines that show strong Th2-bias (Zhu et al., 2021). The balanced immune response was also uniformly recapitulated in a dose response experiment. Nearly the same levels of Th1 and Th2 antibody responses were elicited with the medium-dose as with the high-dose, while the levels were lower (5-25-fold) with the low-dose or single-dose antigen (Figure S1).

Intriguingly, the T4-CoV-2 vaccine induced high levels of spike-specific serum IgA antibodies when administered by either the i.m. or the i.n. route (Figures 2I and 2J). This is notable because IgA stimulation is not commonly observed in traditional vaccines including the current COVID-19 vaccines. For example, the adenovirus-based vaccines do not elicit significant spike-specific serum IgA titers when injected by the i.m. route (Hassan et al., 2020). Elicitation of serum IgA is considered desirable for an effective COVID-19 vaccine because IgA antibodies are reported to have anti-inflammatory activity and are more potent than IgG in neutralizing SARS-CoV-2 virus during the early phase of infection (Sterlin et al., 2021b).

#### Virus neutralizing antibodies

To further analyze humoral immunity, the virus neutralizing activity of the elicited antibodies was determined by Vero E6 cell cytopathic assay using SARS-CoV-2 WA-1/2020 ancestral strain in the US (Harcourt et al., 2020). As shown in Figure S2A, the T4-CoV-2 vaccine induced strong neutralizing activity in sera of all immunized mice. Significantly higher neutralizing antibody titers were detected in mice immunized i.m. with 2 doses of the T4-CoV-2 vaccine than with a single dose immunization (Figure S2A). Importantly, a higher neutralizing antibody titer (3-fold) was induced by i.n. vaccination when compared to i.m. route of high-dose immunization (Figure S2A).

It is well known that Beta and Delta variants escape vaccine-induced immune responses (Mistry et al., 2022). Intriguingly, the T4-CoV-2 vaccine elicited comparable virus neutralizing activities to WA-1/2020, Beta (B.1.351), and Delta (B.1.617.2) VOCs (Figure 1K). Additionally, ~3-fold higher neutralizing antibody titer against SARS-CoV-2 and its VOCs was elicited by i.n. vaccination of mice when compared to i.m. route of immunization, while no detectable neutralizing activity was detected in T4 vector or PBS control groups (Figure 1K).

#### Cell-mediated immunity

To evaluate cellular immune responses, splenocytes were harvested from mice on day 26 after the boost (Figure 1B). Antigen-specific CD8^+^ and CD4^+^ T cells were identified after *ex vivo* restimulation with either S-trimer (Figures 2L and 2M; Figures S2B and S2C) or with SARS-CoV-2 peptides spanning the S- and NP-proteins (Figures S2D and S2E). The samples were then analyzed by intracellular staining of accumulated cytokines and flow cytometry. The percentages of CD8^+^ and CD4^+^ T cells positive for interferon (IFN)-γ, tumor necrosis factor (TNF)-α, or interleukin 17A (IL-17A) were elevated in T4-CoV-2 immunized mice as compared to the T4 vector control group irrespective of the immunization routes and the virus-specific stimulants used (Figures 2L and 2M; Figures S2B to S2E).

IFNγ is a predominant cytokine secreted by effector CD8^+^ T cells, Th1 CD4^+^ T cells, and NK cells (Castro et al., 2018). More specifically, with re-stimulation of splenocytes using S protein, significant levels of IFNγ^+^ CD8^+^ cells, which play a critical role in SARS-CoV-2 viral clearance, were observed in i.n.-immunized mice (Figure 2L). Additionally, significantly elevated percentages of CD4^+^ T cells producing IFNγ were detected in the i.n. group in comparison to the i.m. group of vaccinated mice (Figure 2M, Figure S2C). These data indicated an enhanced Th1-mediated immunity induced by i.n. administration of the vaccine. Of note, we did not observe significant differences between i.n. and i.m. routes of immunization regarding either the IFNγ^+^ CD8^+^ cells or the IFNγ^+^ CD4^+^ cells when restimulated with S- and NP-peptides (Figures S2D and S2E). Probably the conformational epitopes in S- and NP-proteins could contribute to these differences of higher IFNγ levels in the i.n. group of animals. The robust T cell cytokine responses paralleled greater T cell proliferation in both i.n. and i.m. immunized groups of animals as compared to the T4 vector control group (Figures S2F and S2G).

Additionally, representative Th1 and Th2 cytokines in cell supernatants of the splenocytes were analyzed by Bio-Plex platform. Both routes of immunization triggered increased production of Th1 cytokines (IFNγ, IL-2, TNFα, and IL12-p70) (Figures 2N to 2Q; Figures S2H to S2K) and Th2 cytokines (IL-4, IL-5, IL-10, and IL-13) (Figures 2R to 2U; Figures S2L to S2N) compared to controls when splenocytes were stimulated with S-trimer (Figures 2N to 2U) or S- and NP-peptides (Figures S2H to S2N). Increases in Th1 and Th2 cytokine levels by T4-CoV-2 immunization were consistent with induction of balanced Th1 and Th2 antibodies and cellular immune responses, as described above. Importantly, the levels of the main Th1 cytokines, including IFNγ, IL-2, and TNFα, were significantly higher in animals immunized by the i.n. route than those in mice immunized by the i.m. route (Figures 2N to 2P; Figures S2H to S2J). These data indicated that T4-CoV-2 i.n. immunization most likely produced more Th1-biased immune responses. The vaccine-associated enhanced respiratory disease has not usually occurred when strong Th1 cell responses are induced. Therefore, considering that COVID-19 vaccine designs developed to date have attempted to elicit either a Th1-biased or a Th1/Th2-balanced cell response (Sadarangani et al., 2021; Sette and Crotty, 2021), the T4-CoV-2 vaccine generated the desirable responses.

### Needle-free T4-CoV-2 vaccination elicits robust mucosal immune responses

It is generally recognized that i.n. vaccination leads to higher levels of sIgA antibodies at the mucosal surface with lower systemic IgG antibodies and cellular immune responses, while the opposite is true for i.m. vaccination (Krammer, 2020; Macpherson et al., 2008; Su et al., 2016; Tiboni et al., 2021). Remarkably, however, i.n. T4-CoV-2 vaccination induced higher systemic as well as mucosal immune responses (Figures 2 and 3). This appears to be a distinctive feature of the T4 nanoparticle vaccine.

**Figure 3.**
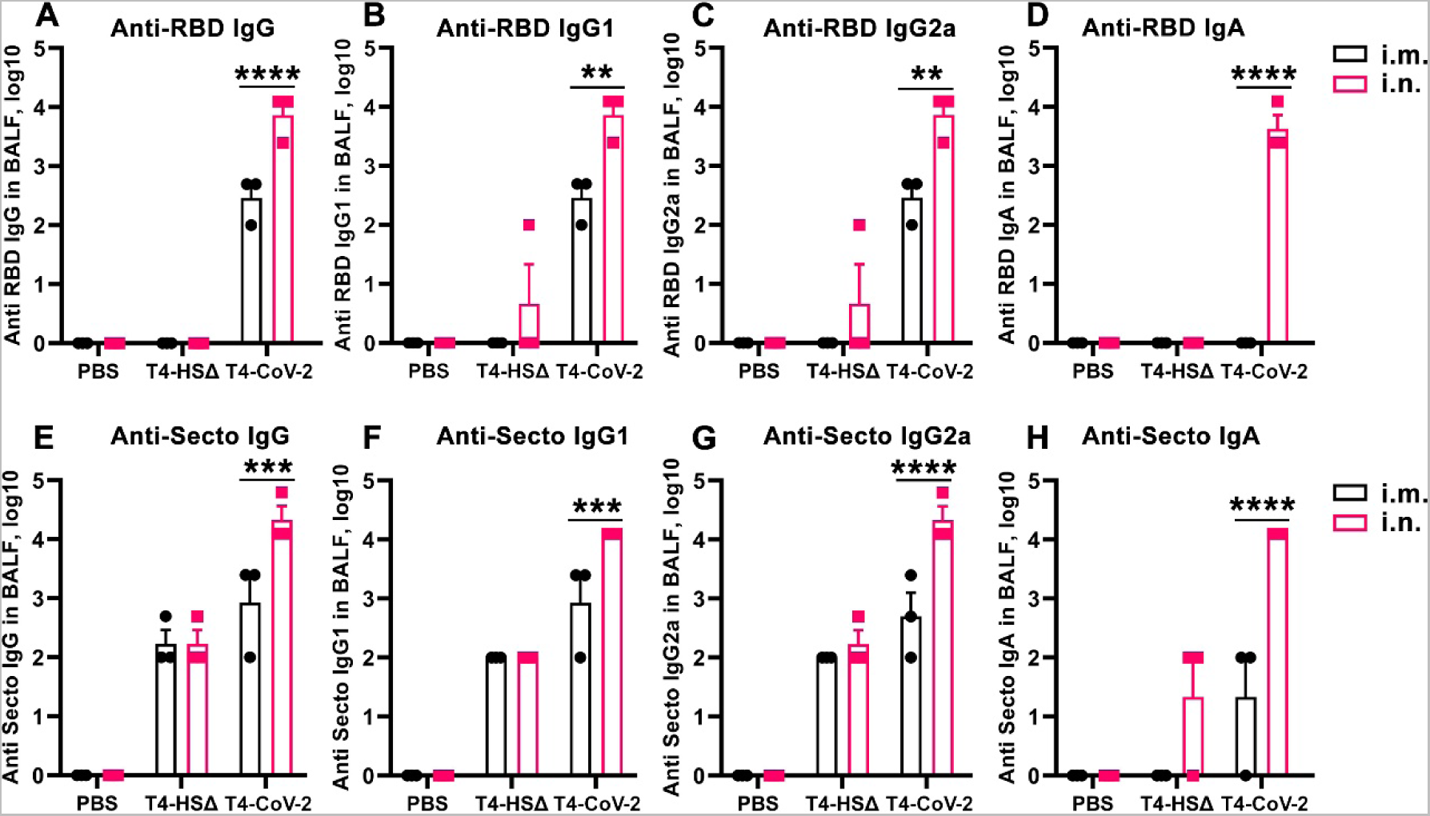
Intranasal immunization with T4-CoV-2 vaccine induced robust mucosal immune responses. The reciprocal endpoint antibody titers in BALF of anti-RBD IgG **(A)**, anti-RBD IgG1 **(B)**, anti-RBD IgG2a **(C)**, anti-RBD IgA **(D)**, anti-Secto IgG **(E)**, anti-Secto IgG1 **(F)**, anti-Secto IgG2a **(G)**, and anti-Secto IgA **(H)** are shown. Data represent mean ± SEM. Data are from 3 pooled independent experiments (n = 12 for T4-CoV-2, n = 10 for T4-HSΔ, and n = 5 for PBS). The titers between i.m. and i.n. route were compared and statistically analyzed by two-way ANOVA test; **P<0.01, ***P<0.001, ****P<0.0001.

Indeed, the needle-free T4-CoV-2 vaccine induced robust mucosal IgG and sIgA responses. These anti-RBD or anti-Spike antibody titers were determined in bronchoalveolar lavage fluid (BALF) samples of vaccinated mice after the booster dose (Figure 3). Intranasally administered vaccine elicited ~25-fold higher IgG antibody levels in BALF compared to when animals were vaccinated by the i.m. route (Figures 3A and 3E), which also included both the Th1-biased IgG2a and Th2-biased IgG1 subtype antibodies in a balanced manner (Figures 3B, 3C, 3F, and 3G).

The sIgA antibodies play a critical role in protecting mucosal surfaces against pathogens by blocking their attachment and/or entry of viruses transmitted through the respiratory tract. Thus, most significantly, high titers of mucosal sIgA antibodies were elicited by i.n. vaccination (Figures 3D and 3H), in addition to high levels of systemic immune responses as described above (Figure 2). In contrast, i.m. immunization failed to produce sIgA, which is not unexpected (Figures 3D and 3H). Since IgA antibodies are dimeric, they might have stronger SARS-CoV-2 viral neutralization activity, and therefore, could confer protection at the site of exposure because mucosal surfaces of the respiratory tract, including the nasal regions and lung epithelial cells, are the major targets for SARS-CoV-2 infection (Figure 1Ee) (Asahi-Ozaki et al., 2004; Lapuente et al., 2021; Renegar et al., 2004).

### Needle-free T4-CoV-2 vaccine provides complete protection and apparent sterilizing immunity against SARS-CoV-2 challenge

#### Animal challenge

BALB/c mice were challenged with the mouse-adapted SARS-CoV-2 strain (MA10) (Leist et al., 2020) (Figure 2B). As shown in Figures 4A to 4D, the control animals that received the T4 vector exhibited a rapid weight loss soon after infection, with a maximum decrease on days 2-4 (Figures 4A and 4B). On the other hand, mice immunized with the T4-CoV-2 vaccine by either of the two immunization routes showed modest-to-no weight loss over the course of 7 days after challenge. However, the data were more impressive after i.n. immunization.

**Figure 4.**
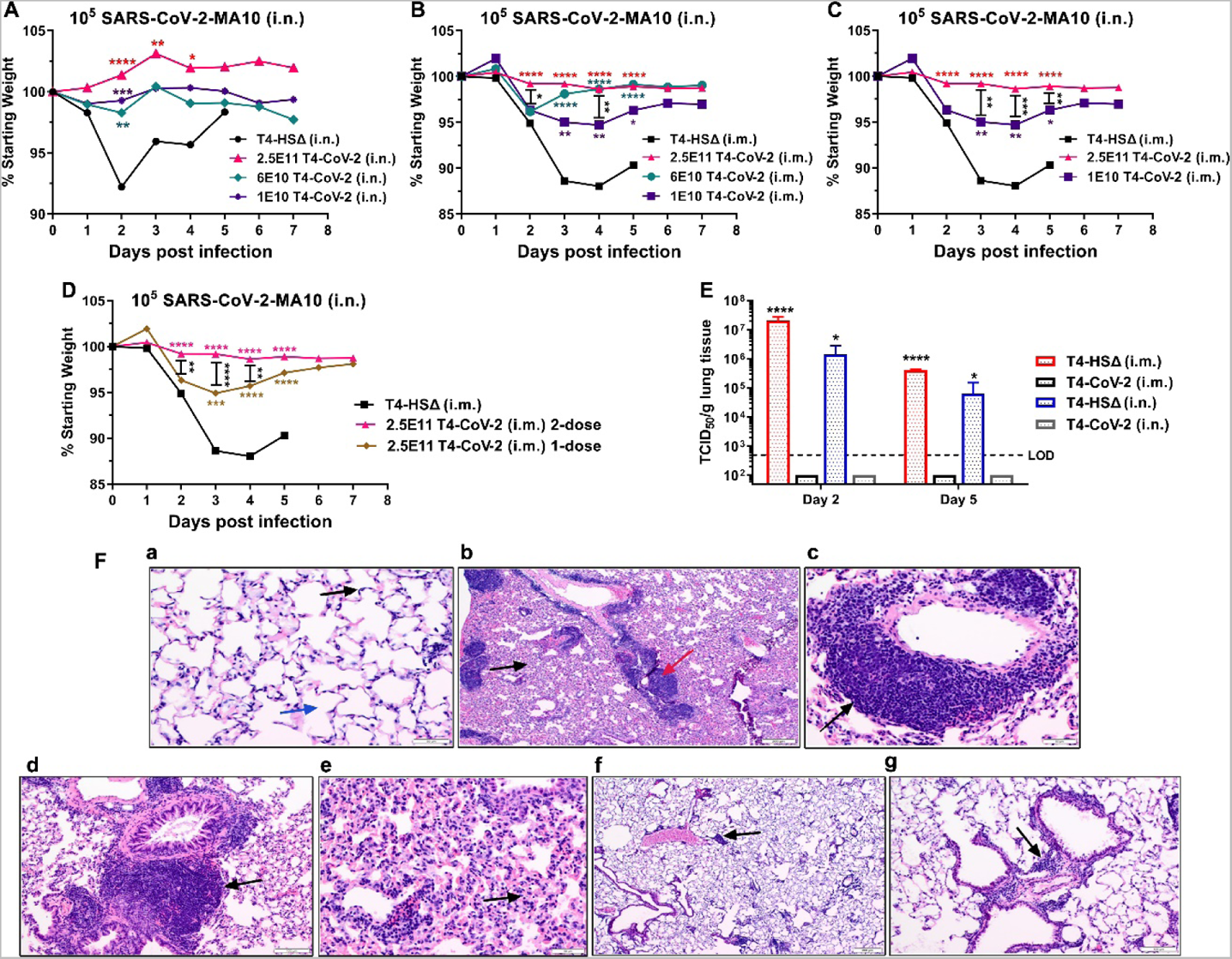
Needle-free T4-CoV-2 vaccination provided complete protection against SARS-CoV-2 challenge.) Percentage starting body weight of i.n. (A) and i.m. (B to D) immunized mice at days after intranasal challenge with SARS-CoV-2 MA10. **(E)** Viral burden (TCID50/g lung tissue) in the lungs at 2 days and 5 days post-SARS-CoV-2 MA10 infection. T4-CoV-2 immunization was compared with vector control in either i.m. or i.n. groups. Dotted lines indicate the limit of detection (LOD) of the assay. **(F)** Histopathological analysis of lung tissues from the vector control and T4-CoV-2 i.n. immunized and challenged mice. Representative photomicrographs from each group are shown. **a.** Medium power view of normal lung with delicate alveolar septa (black arrow) and distinct alveolar spaces (blue arrow) (200X). **b.** Low power view of lungs of the challenged control mice with prominent inflammatory infiltrates of bronchovascular bundles (red arrow), as well as interstitial involvement (black arrow; 40X). **c.** Medium power view of mononuclear inflammatory infiltrates around pulmonary vessel (black arrow; 200X) in the challenged control mice. **d.** Medium power view of mononuclear cell infiltrate around bronchovascular bundle (black arrow; 100X) in the challenged control mice. **e.** Medium power view of distal airways with evidence of interstitial inflammation in alveolar septa (black arrow) in the challenged control mice. **f.** Low power view of lung with mild and patchy inflammatory infiltrate of bronchovascular bundles (black arrow) in the challenged T4-CoV-2 immunized mice. Alveolar spaces and interstitium appear normal (40X). **g.** Medium power view of inflammatory infiltrate around bronchovascular bundle (black arrow; 100X) in the challenged T4-CoV-2 vaccinated mice. A-D, two-way ANOVA with Tukey’s post hoc test to compare multiple groups; E, one-way ANOVA with Tukey’s post hoc test (i.m.) and Mann-Whitney U test (i.n.) (n = 2-5). *P < 0.05; **P < 0.01, ***P < 0.001, ****P<0.0001.

More specifically, the weight loss curves among the high, the medium, and the low dose groups of i.n. vaccination were almost similar statistically. Compared to the T4 vector control, a much-reduced loss in body weights were noted on day 2 post infection (p.i.) in all of the T4-CoV-2 vaccinated groups of mice with subsequent minimal and statistically insignificant fluctuations in body weight changes until day 7 (Figure 4A).

In i.m.-immunized groups, a similar comparison showed statistically significant differences on different days (Figure 4B). Significantly less efficacy of the vaccine was apparent when the number of phage particles was reduced from 2.5 × 10^11^ to 1 × 10^10^ between days 3-5 p.i. (Figure 4C). Similarly, significantly more weight loss was noticed in mice i.m. immunized with one dose of the T4-CoV-2 vaccine as compared to those receiving two doses on days 2-4 p.i. (Figure 4D). These data were consistent with the lower levels of immune responses elicited by i.m. vaccination when compared to i.n. vaccination.

#### Viral load

To further assess protective efficacy in the lungs, the infectious virus load was determined by plaque assay on days 2 and 5 p.i., the peak period of viral burden in this model. As shown in Figure 4E, no infectious SARS-CoV-2 virus could be detected in the lungs of mice immunized with the T4-CoV-2 vaccine (2.5 × 10^11^ phage particles) by either the i.m. or the i.n. route. Quite the opposite, very high levels of virus, ~10^5^ to 10^7^ TCID_50_/g (Tissue culture infectious disease [TCID]), were present on day 2 of the control mice, which decreased substantially on day 5 p.i., at which time the survived animals began to recover from infection. This indicates that the vaccine might be inducing sterilizing immunity, hence minimizing live virus shedding. This is consistent with the induction of strong mucosal immunity as evident from S-specific IgG and sIgA responses in the lungs of i.n.-vaccinated mice. However, even the i.m.-vaccinated mice showed sterilizing immunity suggesting that the relatively low levels of mucosal immunity due to S-specific IgG in lungs combined with the strong CD8^+^ cytotoxic T cells might be sufficient to clear the virus-infected cells.

#### Histopathology

The lung tissues obtained from the control and immunized mice were subjected to H&E (hematoxylin and eosin) and MOVAT staining for histopathological analysis. The analysis was performed based on three parameters: mononuclear inflammatory infiltrate around bronchovascular bundles, interstitial inflammation, and alveolar exudate/hemorrhage.

As shown in Figure 4F, the uninfected normal lungs had delicate alveolar septa (black arrow) and distinct alveolar spaces (blue arrow) with no evidence of inflammation, hemorrhage, exudates, or transudates (panel a, 200x). On the other hand, prominent inflammatory infiltrates of bronchovascular bundles (red arrow) as well as interstitial involvement (black arrow) were noticed in the T4-vector control mice (i.n. immunized) during virus infection (panel b, 40x). More specifically, mononuclear inflammatory infiltrates were noticed around pulmonary vessel (black arrow, panel c, 200x) and bronchovascular bundle (black arrow, panel d, 100x). Distal airways with interstitial inflammation in alveolar septa (black arrow, panel e) were evident. In addition, alveolar hemorrhage was also observed in other areas of the lungs.

As for the T4-CoV-2 i.n. immunized mice, only mild and patchy inflammatory infiltrate of bronchovascular bundles (black arrow, panels f & g, 40x and 100x, respectively) were noted after infection and the alveolar spaces and interstitium appeared normal (panels f and g). Such minimal infiltrates in the lungs were also observed in SARS-CoV-2 mRNA and adenovirus vaccines (Corbett et al., 2020; Hassan et al., 2020). Overall, the combined scores based on the above three parameters were 6.2±1.3 for the T4 vector control and 4.4±1.1 for the T4-CoV-2 vaccine i.n. immunized animals (p = 0.01) when combined data on tissues after 2 and 5 days of challenge were analyzed.

Collectively, T4-CoV-2 vaccination by either route completely protected mice against SARS-CoV-2 challenge with no significant infectious virus detectable and marked attenuation of the inflammatory response in the lungs. These data indicated that the T4-CoV-2 vaccine was effective in clearing the virus and potentially could block transmission of SARS-CoV-2.

### A Beta-variant needle-free T4-CoV-2 vaccine stimulates strong mucosal, humoral, and cellular immune responses in human ACE2 (hACE2) transgenic mice

To determine if the robust and diverse immune responses elicited by the T4-CoV-2 vaccine, especially the mucosal responses, could be recapitulated in highly susceptible hACE2 knock-in mice, we conducted an independent study. Additionally, we constructed a beta-variant spike trimer (Secto-β) (without any affinity tags) for vaccination as this was a dominant strain at the time of the study causing a major second wave in South Africa and across the globe (Tegally et al., 2021). Secto-β contained four critical mutations (K417N, E484K, N501Y, and D614G) that conferred enhanced transmissibility and lethality, and also partial escape from vaccine-induced immunity (Ahmad, 2021) (Figure S3A). The Secto-β variant trimer conjugated to T4 capsid as efficiently as the WT S-trimer through the Spytag-SpyCatcher system (Zhu et al., 2021) (Figure S3B). In addition, the T4-CoV-2-β vaccine also contained ~100 copies of NP protein packaged inside the capsid (Figure S3C). Five-week-old hACE2 AC70 mice were immunized with this vaccine using the same prime-boost regimen (Figure 5A and 5B) at a high dose (~2.5 × 10^11^ phage particles decorated with 20 µg of variant Secto-β).

**Figure 5.**
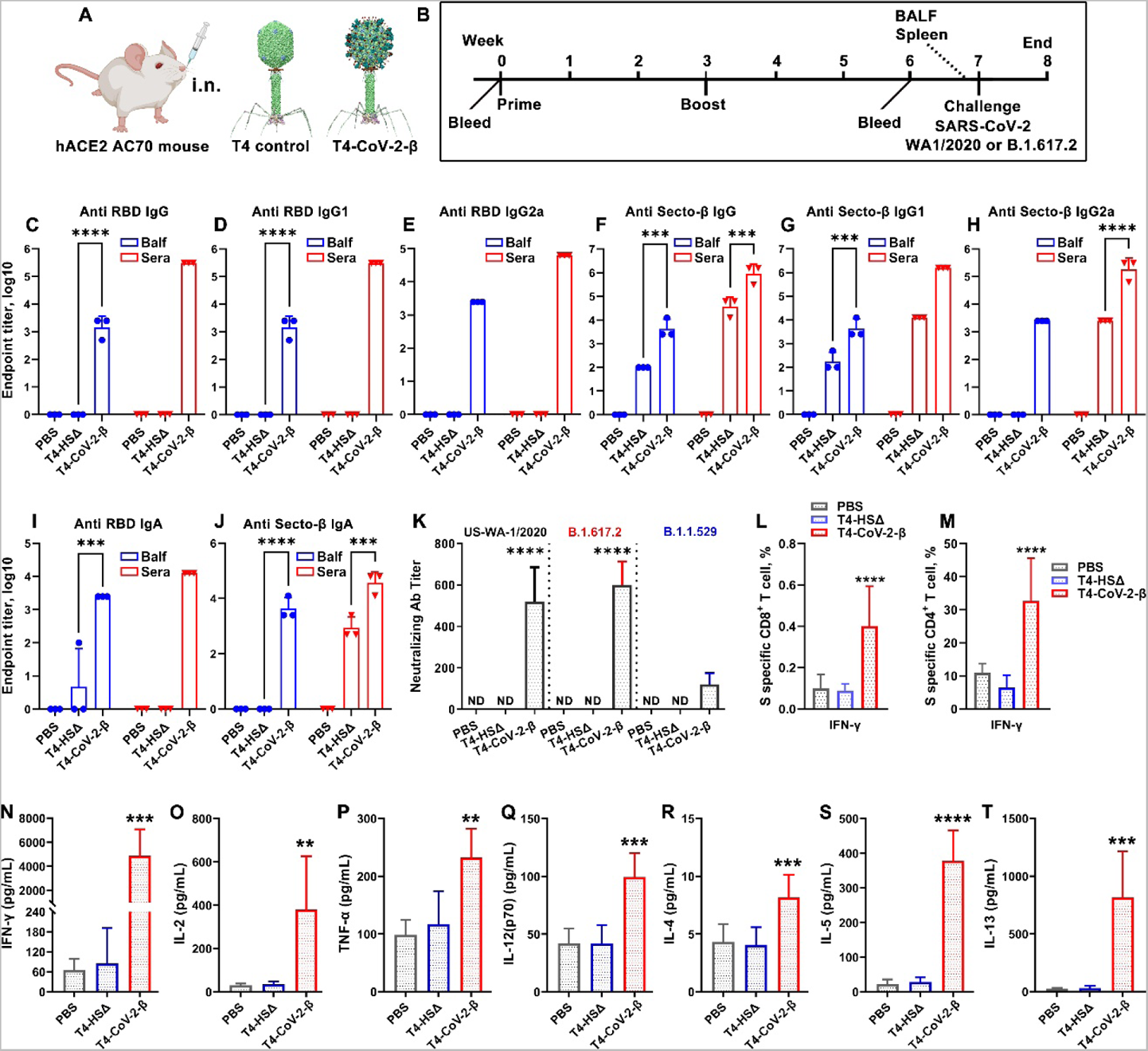
Intranasal T4-CoV-2-β vaccination stimulated robust mucosal and systemic humoral and cellular immune responses in human ACE2 (hACE2) transgenic mice. **(A)** Schematic of intranasal mouse vaccination with T4-HSΔ control or T4-CoV-2-β vaccine. **(B)** Scheme for vaccination and challenge. **(C to J)** Antibody responses in sera (red) and BALF (blue) of immunized mice on day 21 after the boost. ELISA assay was applied to determine reciprocal endpoint antibody titers of anti-RBD IgG (C), anti-RBD IgG1 (D), anti-RBD IgG2a (E), anti-Secto-β IgG (F), anti-Secto-β IgG1 (G), anti-Secto-β IgG2a (H), anti-RBD IgA (I), and anti-Secto-β IgA (J). **(K)** Neutralizing antibody titers in sera were determined by Vero E6 cell cytopathic assay using WA-1/2020, B.1.617.2 (Delta), and B.1.1.529 (Omicron) strains. **(L and M)** Cellular immune responses after stimulation with Secto-β protein. Percentage of IFNγ^+^ CD8^+^ (L) and IFNγ^+^ CD4^+^ (M) cells were plotted. **(N to T)** Splenocyte cytokine responses to Secto-β protein stimulation in immunized hACE-2 transgenic mice. Representative Th1 (N to Q) and Th2 (R to T) cytokines are shown. C to M, two-way (C to J, L to M) or one-way ANOVA (K) with Tukey post hoc test; N to T, nonparametric Student’s t test. Data are from 3 pooled independent experiments (n = 15 for T4-HSΔ and PBS sera analysis, n = 21 for T4-CoV-2-β sera analysis, and n = 5 for BALF analysis). Data are representative of two (K) or five biological replicates (L to T). **P < 0.01, ***P<0.001, ****P<0.0001.

#### Humoral immune responses

Similar to the binding antibody titers in BALB/c mice (Figures 2 and 3), i.n. immunization with T4-CoV-2-β induced high levels of Spike- and RBD-specific IgG and IgA in sera of hACE2-transgenic mice (Figures 5C to 5J), suggesting a strong systemic humoral immune response. In addition, moderate NP-specific IgG antibodies were also elicited in the T4-CoV-2-β immunized mice (Figure S3D). Furthermore, high levels of Spike- and RBD-specific IgG and sIgA antibodies were also present in BALF of T4-CoV-2-β vaccinated mice indicating an equally robust mucosal immune response (Figures 5C to 5J, Figures S4A to S4D). Finally, balanced Th1 and Th2 antibody responses were induced both in sera and BALF, in T4-CoV-2-β immunized mice (Figures 5D, 5E, 5G, and 5H; Figures S4B and S4C). There was no significant difference in binding antibody titers between Secto and Secto-β as the coating antigen (Figures S4E and S4F), probably because they share a large number of the same epitopes. Collectively, consistent with our findings in BALB/c mice, T4-CoV-2-β i.n. vaccination stimulated strong mucosal and systemic humoral immune responses in hACE2-transgenic mice.

Importantly, consistent with the broad-spectrum neutralizing activities in BALB/c mice (Figure 1K), T4-CoV-2-β vaccine elicited comparable virus neutralizing activities to WA-1/2020 and its Delta (B.1.617.2) VOC in hACE2-transgenic mice, while no detectable neutralizing activities were detected in PBS or T4 vector control groups (Figure 5K). Additionally, the Omicron (BA.1) variant emerged in late November of 2021 (near the end of this study) and has the largest number (>30) of mutations within the spike protein described to date. These mutations substantially jeopardized the efficacy of existing COVID-19 vaccines (Edara et al., 2022; Ying et al., 2022), resulting in a major spike in breakthrough infections. Our T4-CoV-2-β vaccinated sera neutralized the Omicron variant (B.1.1.529) but the titers were 6-fold lower when compared to the WA-1/2020 strain (Figure 5K). Interestingly, neutralization of Omicron was comparable to that of WA-1/2020 in BALF (Figure S5A), although the BALF titer appeared lower than that of sera, largely due to dilution of the lung lining fluid.

#### Cell-mediated immune responses

As shown in Figures 5L and 5M, restimulation of splenocytes *ex vivo* with S protein showed a similar pattern of CD8^+^ and CD4^+^ T cell activation in hACE2 mice as with the conventional BALB/c mice (Figures 2L and 2M). The percentages of CD8^+^ and CD4^+^ T cells positive for IFNγ were substantially elevated in T4-CoV-2-β immunized mice as compared to both PBS and T4 vector control groups (Figures 5L and 5M). Interestingly, a much higher percentage of IFNγ positive CD4^+^ T cells was observed in hACE2 mice than those in conventional BALB/c mice, while the percentage of TNFα or IL-17A positive T cells were similar (Figures S5B and S5C). T4-CoV-2-β i.n. immunization developed robust spike-specific CD8 and CD4 T cell responses in hACE2-transgenic mice.

Similarly, both Th1 cytokines (IFNγ, IL-2, TNFα, and IL12-p70) (Figures 5N to 5Q) and Th2 cytokines (IL-4, IL-5, and IL-13) (Figures 5R to 5T) were induced in T4-CoV-2-β-immunized mice compared to the controls when splenocytes were re-treated with the Secto-β trimer. Significantly, very prominent Th1 cytokines IFNγ and IL-2 were produced, indicating Th1-biased cellular immune responses induced by intranasal T4-CoV-2-β vaccine.

### Needle-free T4-CoV-2-Beta vaccine provides complete protection and apparent sterilizing immunity against lethal infection by both the original SARS-CoV-2 and the Delta VOC in hACE2 transgenic mice

#### Animal challenge and viral load

Mice were i.n. challenged with either WA-1/2020 strain or its Delta (B.1.617.2) variant. The highly contagious B1.617.2 shows increased transmissibility compared to the ancestral strain, and studies suggested a high risk of hospitalization compared to the original strain (Liu and Rocklöv, 2021). As shown in Figure 6A, irrespective of the challenge strains, all control animals rapidly lost weight (Figure 6A) and succumbed to infection (Figures 6B and 6C) on day 4-5 post challenge. In contrast, all the T4-CoV-2-β immunized mice only had minimal to no weight loss with a 100% survival rate over 21 days after the challenge. Furthermore, a high viral load in the lungs was observed in all the control animals on day 5 p.i., while no live virus was detected in the lungs of T4-CoV-2-β vaccinated mice (Figure 6D).

**Figure 6.**
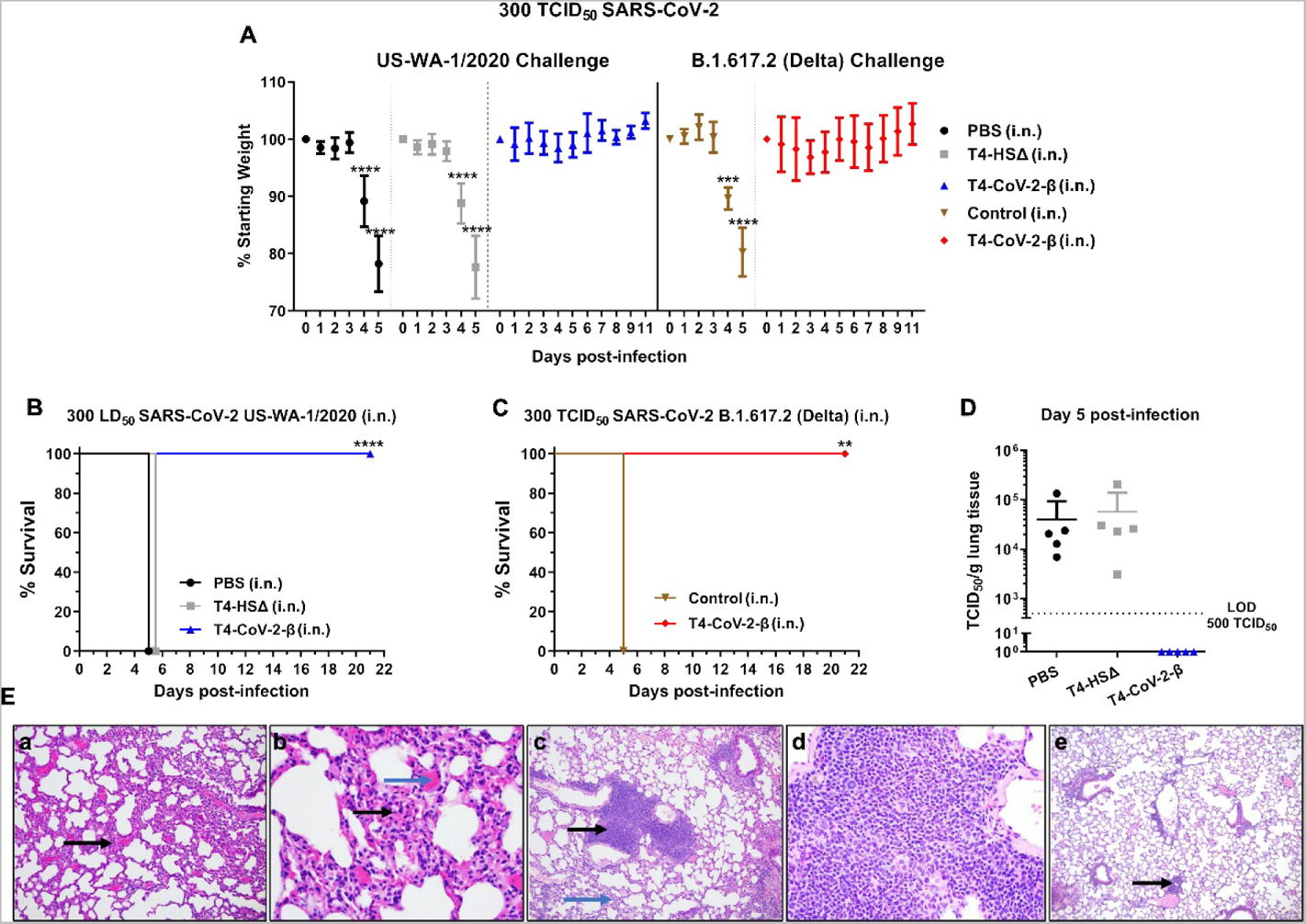
Needle-free T4-CoV-2-β vaccine provided complete protection against lethal infection by ancestral SARS-CoV-2 strain as well as its Delta variant in hACE2 transgenic mice. **(A)** Percentage starting body weight of immunized mice on various at days after i.n. challenge with 300 TCID_50_ of WA-1/2020 strain or its Delta (B.1.617.2) variant. **(B and C)** Survival rate of hACE2 transgenic mice immunized with T4-CoV-2-β or T4-HSΔ vector control against WA-1/2020 strain (B) or its Delta variant (B.1.617.2) (C). **(D)** Viral burden (TCID_50_/g lung tissue) in the lung at 5 days post WA-1/2020 infection. Dotted lines indicate the limit of detection (LOD) of the assays. **(E)** Lung tissues obtained from the control (**a and b**) and T4-CoV-2-β (**c to e**) immunized mice (i.n.) were subjected to H&E (hematoxylin and eosin) and MOVAT staining for histopathological analyses, and representative photomicrographs from each group are shown. A, multiple Student’s t-test using Holm-Sidak method to correct for multiple comparisons (n = 3-10); B to C, Kaplan Meier analysis with log-rank (Mantel-Cox) test (n = 3-10) **P < 0.01, ***P < 0.001, ****P<0.0001.

#### Histopathology

As can be seen from Figure 6E, hACE2 transgenic mice treated with PBS and then challenged with WA-1/2020 strain showed significant interstitial inflammation in alveolar septa (black arrow, panel a, 100x) and alveolar hemorrhage. However, there was no evidence of bronchovascular inflammatory infiltrates on day 5 p.i. At 200x, widening of interstitium with mononuclear inflammatory infiltrates (black arrow) and septal capillary congestion was clearly visible (blue arrow, panel b) in PBS treated and challenged mice.

Based on interstitial inflammation, animals receiving PBS or immunized with T4 vector and then challenged had similar scores of 40±7.1 (PBS group) and 46±18 (T4 vector control group) on day 5 p.i., and the data were not significantly different (p=0.5, Student’s t test). Further, on comparing unvaccinated animals (PBS + vector control groups together) with animals receiving the T4-CoV-2-β vaccine, interstitial inflammation was significantly less in immunized mice (p=0.007, Mann-Whitney rank-sum test, the results were expressed as median, 25%, and 75% with values of 40, 30, and 52.5 for PBS and T4 vector control immunized and challenged mice compared to 20, 20, and 30 for the T4-CoV-2-β vaccinated and challenged animals) on day 5 p.i.

Although T4-CoV-2-β i.n. vaccinated and challenged animals had mild interstitial inflammation (blue arrows, panel c), bronchovascular inflammatory infiltrates (black arrows, panel c, 100x) were clearly visible, not noted in unvaccinated and challenged mice. The bronchovascular infiltrates were mainly composed of lymphocytes and scattered macrophages (200x, panel d). Statistically, mice vaccinated with the T4 vector or T4-CoV-2-β and then challenged had a higher level of bronchovascular infiltrates than PBS treated and infected animals, indicating that T4 phage could increase bronchovascular infiltrates in hACE2 transgenic mice. Importantly, at day 30 p.i., there was no evidence of interstitial pneumonitis and only a mild bronchovascular inflammation (black arrow, panel e, 40x) in T4-CoV-2-β vaccinated and challenged mice. These data indicated almost complete recovery of animals from bronchovascular infiltrates.

Overall, our data indicated immunological responses induced by the vaccine cleared the infection with 100% survival of the animals. T4 vector, like any other vectors, is expected to activate some non-specific and non-damaging immune responses in the host which subside as the vaccine clears from the host.

### The T4-CoV-2 vaccine is stable at ambient temperature

The current mRNA vaccines require sub-freezing temperatures, and the adenovirus-based vaccines require cold temperatures, for storage and distribution. Bacteriophage T4 being a resident of the gut has evolved a stable capsid structure to survive in a hostile environment. Indeed, the T4 phage is stable at extremes of pH and at ambient temperature, properties that are particularly suitable for storage and extending the life of a vaccine (Jończyk et al., 2011).

To determine the stability of the T4-CoV-2-β, the vaccine preparations in PBS were stored at 4°C and room temperature (22°C), and samples were taken at various time points and analyzed for stability and functionality. Stability was assessed by any reduction in the amount of intact spike protein associated with phage (due to dissociation), and/or appearance of any degraded protein fragments (due to nonspecific proteolysis), whereas functionality was assessed by the ability of the displayed S-trimers to bind to hACE2 receptor. The data showed (Figures 7A to 7D) that the T4-CoV-2-β vaccine, by any of these criteria, was completely stable and functional for at least 10-weeks of storage at 4°C or at 22°C. Furthermore, the backbone phage displaying the SpyCatcher domain as part of the hard-wired recombinant phage, i.e., prior to conjugation with S-trimer, also remained completely stable and functional.

**Figure 7.**
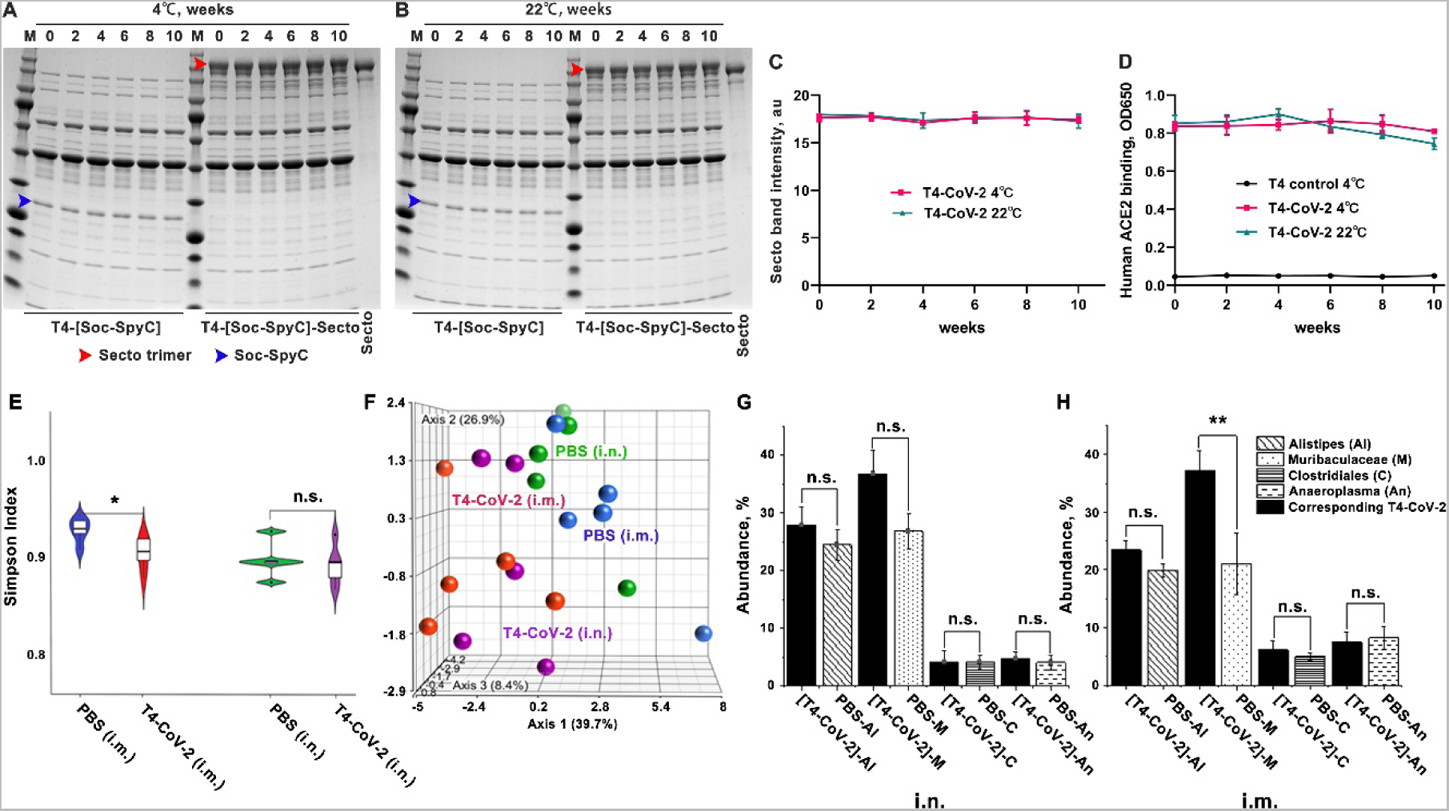
T4-CoV-2 vaccine is stable at ambient temperature and does not influence the microbiome community in mice. **(A and B)** The stability of T4-CoV-2 and T4-(Soc-SpyC) phages for 10-weeks at 4°C (A) or 22°C (B). Samples were taken every two weeks and analyzed for stability by SDS-PAGE. The blue and red arrowheads indicate the bands of Soc-SpyCatcher and covalently conjugated Secto protein, respectively. **(C)** Quantification of the displayed Secto band in T4-CoV-2 vaccine stored at 4°C or 22°C. **(D)** Comparison of binding efficiency of T4–CoV-2 phage to hACE2 receptor after storage at 4°C or 22°C. **(E)** The correlated distribution of the Simpson diversity index of microbiomes from PBS control and T4-CoV-2 vaccinated groups when immunization occurred by the i.n. or the i.m. route. The measure of diversity included number and relative species abundance. **(F)** Summary of individual Euclidian distance as a 3D resemblance matrix of microbial species in the tested groups. **(G and H)** Specific effect of the vaccination of T4-CoV-2 and PBS control on the bacterial genera of the microbiome. The abundance of the gut microbiota following i.n. (G) or i.m. (H) administration of the T4-CoV-2 vaccine and the PBS control are shown.

These data demonstrated stability advantage of the phage T4-CoV-2-β vaccine and coupled with the needle-free i.n. route of administration, this platform provides especially useful features for rapid vaccine distribution during a pandemic (Kim et al., 2021).

### T4-CoV-2 vaccination does not influence the microbiome community

Finally, we determined if T4-CoV-2 vaccination impacted the microbiome community. DNA was extracted from the fecal matter of individual mice (n=5/group) and was sequenced for 16S rRNA gene and analyzed.

#### Violin plot

The violin plot in Figure 7E showed the correlated distribution of the Simpson diversity index of microbiomes in the test groups. The measure of diversity included number and relative species abundance. As noted, the i.n. route of administration did not alter the Simpson diversity of the microbial species recovered from the PBS control versus the T4-CoV-2 vaccine groups of mice, unlike when i.m. route of vaccination was used. These results indicated that i.n. vaccination did not significantly affect the number and relative abundance of the gut microbiota.

#### Principal coordinate analysis (PCoA)

Figure 7F summarized individual Euclidian distance as a 3D resemblance matrix of microbial species. The data indicated that the relative distances based on the number between species during both routes of immunization were similar, but there was a significant difference in species diversity when immunization occurred via the i.m. route (PBS control versus T4-CoV-2 groups). However, this was not the case during i.n. route for immunization as there was a lack of significant differences among species.

#### Specific effect on the bacterial genera of the microbiome

Figures 7G and 7H showed abundance of the gut microbiota. The Tukey mean comparison method between the T4-CoV-2 and PBS groups for the top four genera (*Alistipes*, *Muribaculaceae*, *Clostridiales*, and *Anaeroplasma*) indicated no significant differences in the gut microbiota even though there were few differences in numbers (*e.g.,* for *Alistipes* and *Muribaculaceae*) when vaccine was administered by the i.n. route (Figure 7G). However, a significant difference in the *Muribaculaceae* genus was noted when T4-CoV-2 vaccine was delivered by the i.m. route (Figure 7H). These same differences were observed among the *Bacteroidetes* phylum indicating that i.m. administration of the T4-CoV-2 vaccine had a more significant impact on the gut microbiota. These trends were also reflective upstream of the hierarchy from families to the phylum of the recovered gut microbiota. Post-vaccination microbiota perturbation was previously reported during early microbial and immunological maturation stages in humans (Ruck et al., 2020). Similarly, Chen et al., 2021 identified postvaccination dysbiosis as a significant problem in developing cellular immunity in COVID-19 vaccines, which can be corrected by introducing prebiotics and probiotics oral supplements after vaccination (Chen et al., 2021). However, notably, the T4-based COVID-19 vaccine administered by the i.n. route seemed to circumvent this effect on the microbiota.

## DISCUSSION

A next-generation COVID-19 vaccine that would elicit local mucosal responses in addition to strong systemic immunity is most desired to control SARS-CoV-2 infections and in general, any mucosally transmitted infection (Alu et al., 2022; Borges et al., 2010; Lavelle and Ward, 2021). This is particularly relevant at this stage of the COVID-19 pandemic, in view of the current evolutionary trajectory of the virus selecting highly transmissible variants such as Omicron BA.1 and BA.2.

The sticky mucous layers in the nasal epithelia present barriers to pathogens and possibly interfere with the ability of vaccines to access and activate the mucosal immune system. This may account for poor immunogenicity of most injectable vaccines when administered intranasally (Focosi et al., 2022; Tiboni et al., 2021). At present, of 195 COVID-19 vaccine candidates in clinical trials, only fourteen are intranasal vaccines. Most of them are based on engineered live viruses that can efficiently infect human cells and intracellularly express spike or RBD antigens from the delivered genes. These include human or chimpanzee adenoviruses (Hassan et al., 2020; Hassan et al., 2021; King et al., 2020; Lapuente et al., 2021; van Doremalen et al., 2021a), live-attenuated influenza virus (An et al., 2021; Liu et al., 2021), live-attenuated Newcastle Disease Virus (Park et al., 2021; Sun et al., 2021), and lentivirus (Ku et al., 2021). However, these eukaryotic viral vaccines still pose a safety concern, pre-existing immune responses, and a risk, albeit very low, of reversion.

Our studies established a prokaryotic, noninfectious, bacteriophage T4 mucosal vaccine delivery platform that can be engineered to generate stable, needle- and adjuvant-free, multicomponent vaccines against COVID-19 or any emerging and pandemic pathogen. Presence of ~17-nm long Hoc fibers on T4 capsid surface that could interact with mucin glycoproteins and S-trimers binding to ACE2 receptors provide distinct advantages for intranasal delivery and presentation to host’s mucosal immune system. Indeed, a series of datasets demonstrate that the T4-CoV-2 nanoparticle vaccine containing arrays of ~100 copies of S-trimers on T4 capsid exterior and ~100 copies of NP packaged in its interior when administered to mice intranasally stimulated all arms of the immune system, including strong mucosal immunity that injectable vaccines do not induce.

The immune responses stimulated by the T4 based COVID-19 vaccine were broad and included: Th1 and Th2 derived IgG and IgA antibodies in sera, virus neutralizing antibodies, CD4^+^ helper and effector T cells and CD8^+^ killer T cells, Th1-biased cytokines, and mucosal IgG and sIgA antibodies in BALF. While most of these immune responses were triggered by both i.n. and i.m. routes of vaccine administration, the stimulation was considerably stronger by i.n. immunization. Consistently, weight losses following challenge were significantly lower in the i.n. vaccinated mice than the i.m. mice, although both routes induced apparent sterilizing immunity showing no virus load in the lungs of vaccinated mice. Remarkably, however, the mucosal sIgA in BALF was stimulated only by i.m. vaccination. The sIgA is supposed to be effective at the entry point by interfering with virus acquisition and at the exit point by clearing the invaded pathogen (Figure 1) (Sterlin et al., 2021a; Wang et al., 2021). This pattern of broad responses was consistently observed for both WT as well as the Beta-variant S-trimers and in conventional BALB/c mice as well as hACE2 transgenic mice. The evidence, thus, is compelling to suggest that vaccine-induced mucosal immunity is a prominent feature of the needle-free bacteriophage T4 nanoparticle vaccine, which could be further exploited for designing vaccines against other respiratory infections (Excler et al., 2021).

Strikingly, the T4-CoV-2 vaccine induced similar levels of serum virus neutralizing antibody titers against the ancestral WA-1/2020 strain and its two VOCs (B.1.135 Beta and B.1.617.2 Delta) which can significantly escape immune responses by the existing mRNA or adenovirus vaccines (Kroidl et al., 2021; Mlcochova et al., 2021; Zinatizadeh et al., 2022). Consistently, our vaccine protected mice from challenge by both the WA-1/2020 strain and its Delta (B.1.617.2) variant, considered thus far the most lethal strain. Additionally, the T4-CoV-2 vaccine also induced significant but somewhat diminished neutralizing antibody titers against the Omicron variant which has the greatest number of mutations and immune-escaping capacity reported to date (Ying et al., 2022). Importantly, similar levels of neutralizing antibody titers were measured in BALF against both WA-1/2020 isolate and its Omicron variant. It is intriguing why the neutralization activity induced by T4-CoV-2 vaccination is so broad. One possible reason might be the presence of high levels of sIgA in sera and in BALF, which is reported to be more potent than IgG in neutralizing SARS-CoV-2 virus (Sterlin et al., 2021b).

Notably, the T4-CoV2 nanoparticle vaccine is also a potent inducer of cellular immunity. Our studies demonstrated that both routes of immunization (i.n. and i.m.) induced the enhanced release of pro-inflammatory/anti-inflammatory as well as Th1/Th2 cytokines in BALB/c and hACE2 transgenic mice. Interestingly, i.n. route of immunization induced greater cellular responses, especially Th1, compared to i.m. route of vaccination. Th1 cells and cytotoxic T lymphocytes are primarily responsible for host defense against viral infections, and the role of Th2 cells in recruiting different types of innate immune cells to kill invading pathogens is also well documented (Sallusto, 2016). A Th1 cell-biased response or balanced Th1 /Th2 cell response has also been reported by others upon immunization of mice, hamsters, and macaques with effective COVID-19 vaccines (Bos et al., 2020; Corbett et al., 2020; DiPiazza et al., 2021; Kalnin et al., 2021; Sadarangani et al., 2021; Vogel et al., 2020; (Zhang et al., 2022). Therefore, a combination of producing neutralizing antibodies and activation of antigen-specific T cells may act in concert to control SARS-CoV-2 infection in our mouse models.

Additionally, we also observed Th17 immune responses elicited by T4-COVID vaccine. Th17 cells are being recognized as an important T helper subset for immune-mediated protection, and unbalanced Th17 responses are implicated in the pathogenesis of several autoimmune and allergic disorders (Tesmer et al., 2008). Involvement of IL-17 in priming enhanced chemokine and G-CSF production in the lung during bacterial pneumonia and its ability to promote antimicrobial responses against pathogens of viral, bacterial, parasitic, and fungal etiology has been reported (Anipindi et al., 2019; Guo et al., 2011; Ma et al.). For example, mucosal delivery of *M. tuberculosis* subunit vaccine has been shown to provide IL-17 dependent protection of mice against pulmonary tuberculosis compared to when the vaccine was delivered by the parenteral route (Counoupas et al., 2020). Since the T4-COVID vaccine provided complete protection to mice with much reduced histopathological lesions, our data support the notion that a delicate balance of Th1/Th2/Th17 and mucosal immune responses were critical in developing effective COVID-19 vaccines.

The T4-CoV-2 vaccine is a safe and stable vaccine. A noninfectious phage T4-CoV-2 vaccine with no tropism to human cells and no use of adjuvants or chemical stimulants represent significant advantages. In fact, our previous studies showed that adding adjuvants such as alum or liposomes did not further enhance the levels of immune responses (Rao et al., 2011a; Zhu et al., 2021). Microbiome analyses showed no significant changes in the microbiome diversity in mice vaccinated with the T4-CoV-2 vaccine. In human clinical trials and hundreds of T4 phage vaccine immunizations over the years involving mice, rats, rabbits, and macaque animal models and diverse antigens such as anthrax, plague, and HIV did not identify any significant side effects (Li et al., 2021; Rao et al., 2011b; Tao et al., 2013; Tao et al., 2018a; Zhu et al., 2019). Furthermore, the T4 phage is one of the most stable virus scaffolds known (Jończyk et al., 2011) and our stability studies showed that the T4-CoV-2 vaccine was completely stable at ambient temperature for at least 10-weeks. Therefore, the T4 vaccine that requires no cold chain provides an excellent alternative for global distribution and vaccination of still unvaccinated populations across the world.

Additionally, the T4-CoV-2 vaccine is a strong candidate as an effective booster vaccine. Before this pandemic ends, an additional booster will likely be needed to protect the global population from emerging variants. None of the licensed vaccines used worldwide are needle-free or generate significant mucosal responses, which are critically important for minimizing person-to-person transmission. The T4-CoV-2 vaccine that can boost not only the antibody and T cell immune responses but also induce strong mucosal immunity would be the most beneficial one. Furthermore, more than a billion vaccinations across the globe received the adenovirus-based vaccines, which also stimulate strong anti-vector responses. This pre-existing immunity, particularly the adenovirus capsid neutralizing antibodies, limit the effectiveness of another booster dose using the same vaccine, particularly in the elderly (Chevillard et al., 2022; Lanzi et al., 2011), because vaccine delivery requires efficient infection of human cells which would be compromised by immune clearance (Dicks et al., 2022). Since there is no significant preexisting immunity in humans for T4 (Bruttin and Brussow, 2005), the T4-CoV-2 vaccine would be an excellent alternative to boost more than a billion people who already received the adenoviral vaccines.

In conclusion, we have established a bacteriophage T4-based, protein vaccine platform, complementing the current mRNA and DNA vaccine platforms but with certain advantages in terms of route of administration, engineerability, breadth of immune responses, mucosal immunity, and vaccine stability. In particular, broad virus neutralization activity, both systemic and mucosal, T cell immunity, complete protection, and apparent sterilizing immunity, all induced by the same vaccine mean that the T4-CoV-2 vaccine might be able to block viral entry (host’s viral acquisition) and viral exit (host’s viral shedding), minimizing person to person viral transmission. However, additional studies in animal models (hamsters and macaques), Phase 1 human clinical trials, and GMP manufacturing processes are needed to translate the vaccine into mass production and global distribution. These efforts are currently underway and crucial as more than 10 billion doses of the vaccines are needed across the globe, particularly in middle-to-low-income countries where the affordability of the current vaccines is a big concern due to cost.

## METHODS

### T4 bacteriophages and SARS-CoV-2 strains

The T4-CoV-2 vaccine is a recombinant T4 phage displaying 100 copies of prefusion-stabilized SARS-CoV-2 spike protein ectodomain trimers (S-trimers) on the surface of 120 × 86 nm phage capsid. It also harbors SARS-CoV-2 nucleocapsid protein (NP) packaged in its core and a 12-amino acid (aa) peptide of the putative external domain of E protein (Ee) on the capsid surface. The S-trimers were displayed through interaction with the small outer capsid protein (Soc) which is attached to EXPiCHO-expressed S-trimers via SpyCatcher-SpyTag conjugation. The Ee peptide was attached through fusion to the highly antigenic outer capsid protein (Hoc). The NP, Ee, and SpyCatcher were hard-wired into T4 genome by CRISPR engineering and incorporated into the phage nanoparticle structure during phage infection to make vaccine production easy. The T4 phage without carrying the SARS-CoV-2 components was used as a control for the study.

Mouse adapted SARS-CoV-2 MA10 strain is a gift from Dr. R. Baric, University of North Carolina, Chapel Hill, NC. The first COVID-19 patient isolate SARS-CoV-2 US-WA-1/2020, its Beta (B.1.351), Delta (B.1.617.2), and Omicron (B.1.1.529) VOCs were obtained through CDC and available at the Galveston National Laboratory, UTMB.

### T4 bacteriophage production, purification, display, and stability evaluation

Bacteriophages T4-NP-Ee-(Soc-SpyCatcher) and T4-HSΔ were produced in *E. coli* strain B40 and purified by two rounds of CsCl gradient centrifugation as described previously (Zhu et al., 2021; Zhu et al., 2022). The purified phages were passed through a 0.22-µm filter to remove any minor bacterial contaminants. *In vitro* display of Secto or Secto-β trimer on the T4-NP-Ee-(Soc-SpyCatcher) phage was assessed by co-sedimentation as described previously (Zhu et al., 2021). The phage concentration and copy numbers of displayed antigens were quantified by 4-20% SDS-PAGE. The copy numbers of displayed antigens per capsid were calculated using gp23 (major capsid protein; 930 copies) or gp18 (major tail sheath protein; 138 copies) as internal controls and S-trimer protein standard. The copies of the phage-packaged NP protein were quantified by Western blotting using the commercial rabbit anti-NP antibody (Sino Biological) and NP protein standard (ThermoFisher Scientific) as previously described (Zhu et al., 2021).

For stability evaluation, the T4-CoV-2 vaccine phage (T4-[Soc-SpyC]-Secto), as well as the T4 backbone phage (T4-[Soc-SpyC]), were flash-frozen at −70°C at the time zero, as 100% controls. Two sets of the same phages were stored at 4°C or 22°C, and samples were taken at two-week intervals for ten weeks and were flash-frozen at −70°C. All the samples were thawed and analyzed together for stability and functionality by SDS-PAGE and human ACE2 receptor binding assay as previously described (7). After Coomassie Blue R-250 (Bio-Rad) staining and destaining, the displayed S-trimer protein bands on SDS-PAGE gels were scanned and quantified by ChemiDoc MP imaging system (Bio-Rad) and ImageJ.

### Beta-S-trimer (tag-free) purification

To obtain prefusion-stabilized native-like trimers, Secto or Secto-β trimers were expressed from a recombinant plasmid in ExpiCHO mammalian host cells. The CHO cell growth and Spike recombinant plasmid transfection were performed according to the ExpiCHO expression system User Guide (MAN0014337, ThermoFisher website). The S-trimer expression was under the control of a strong CMV promoter. Cultures were harvested 8 days after transfection by centrifuging the cells at 3000 *g* for 20 min at 4°C. The supernatant (culture medium) containing the expressed S-trimers was recovered and clarified through a 0.22 μm filter (Corning Inc.) for column purification.

The pH of the filtered supernatant (250 ml) was first adjusted to 8 using 1M Tris-HCl, pH 8. Then the supernatant was loaded onto two HiTRAP Q-FF columns connected in tandem and previously equilibrated with wash buffer (100 mM NaCl, 50 mM Tris-HCl, pH 8). The sample was loaded at a flow rate of 1 mL/min, using AKTA Prime-Plus liquid chromatography system (GE Healthcare). The flow-through was collected and diluted with 50 mM Tris-HCl, pH 8 buffer at 1:1 ratio and loaded onto the HiTRAP Q-HP column at a flow rate of 1 mL/min, followed by washing the column with 50 mM NaCl, 50 mM Tris-HCl, pH 8 wash buffer until the absorbance reached the baseline. The trimers were eluted using a 50-600 mM linear gradient of salt in 50 mM Tris-HCl, pH 8 (90 mL total gradient). The peak fractions were run on a 4-20% gradient SDS-PAGE to select fractions with a high ratio of trimers to contaminants. The selected fractions were then pooled and concentrated using 100 kDa filters (Millipore) and loaded to a Hi-Load 16/600 Superdex-200 pg (preparation grade) size-exclusion chromatography column (GE Healthcare) equilibrated with the gel filtration buffer (100 mM NaCl, 50 mM Tris-HCl, pH 8) to further separate the low molecular weight contaminants and obtain purified trimers (ÄKTA FPLC, GE Healthcare). Eluted trimer fractions were assessed on the SDS-PAGE gel to determine the purity and selected fractions were pooled and passed through 0.22 µm filter to sterilize the sample. If needed, the trimers were concentrated using 100 kDa centrifugal filters at 3,500 RPM in a swing bucket rotor. The concentration of the Secto trimers was kept around 1-2 mg/mL. Protein aliquots (1 mL size) were made, flash-frozen in liquid nitrogen, and stored at −80°C until use.

### Mouse immunizations

We followed the recommendations of the NIH for mouse studies (the Guide for the Care and Use of Laboratory Animals). All animal experiments were approved by the Institutional Animal Care and Use Committee of the Catholic University of America (Washington, DC) (Office of Laboratory Animal Welfare assurance number A4431-01) and the University of Texas Medical Branch (Galveston, TX) (Office of Laboratory Animal Welfare assurance number A3314-01). The SARS-CoV-2 virus challenge studies were conducted in the animal BSL-3 (ABSL-3) suite at UTMB. Five-week-old female BALB/c (Jackson Laboratory) or hACE2 transgenic mice AC70 (Taconic Biosciences) were randomly grouped (5-10 animals per group) and allowed to acclimate for 14 days. The phage T4-CoV-2 vaccine was administered by either the i.m. or the i.n. route into the hind legs of mice or naris, respectively. For 2-dose regimen, animals received vaccination at days 0 (prime) and 21 (boost), while for 1-dose regimen, the vaccine was given at day 21. Three different number of phage particles possessing 0.8, 4.8, and 20 µg of S-trimer antigens representing ~ 1.0 × 10^10^, 6 × 10^10^ and 2.5 × 10^11^ phage particles, respectively, were used. Negative control mice received the same volume of PBS or the same amount of T4 control phage (T4 control). Blood was drawn from each animal on day 0 (pre-bleed) and day 42, the isolated sera were stored at −80°C until further use.

### Bronchoalveolar lavage fluids collection

On day 21 after boosting, bronchoalveolar lavage fluids (BALF) were obtained from immunized and control animals by following the protocol as previously described with slight modifications (Van Hoecke et al., 2017). Briefly, the salivary glands were dissected to expose the trachea of euthanized mice (n=5/group). A small incision was made on the ventral face of the trachea and a blunt 26G needle was inserted into the trachea and secured by tying the trachea around the catheter using the floss placed underneath the trachea. An aliquot (600 µL) of PBS loaded into a 1mL syringe was flushed in the lungs and BALF was collected.

### ELISA determination of IgG, IgG subtypes, and IgA antibodies

ELISA plates (Evergreen Scientific) were coated with 100 µL (1 µg/mL) per well of SARS-CoV-2 Secto protein (Sino Biological), SARS-CoV-2 Secto-β protein, SARS-CoV-2 RBD-untagged protein (Sino Biological), SARS-CoV-2 NP (Sino Biological), or SARS-CoV-2 E protein (1 to 75 amino acids) (ThermoFisher Scientific) in coating buffer [0.05 M sodium carbonate-sodium bicarbonate (pH 9.6)] at 4°C for overnight incubation. The plates were washed twice with PBS buffer, followed by blocking with 200 µL per well of PBS (pH 7.4)– 5% BSA (bovine serum albumin) buffer at 37°C for 2 h. Serum and BALF samples were diluted with a 5-fold dilution series beginning with an initial 100-fold dilution in PBS–1% BSA. One hundred microliters of diluted serum or BALF samples were added to each well, and the plates were incubated at 37°C for 1 h. The plates were washed five times with PBST (PBS + 0.05% Tween 20). Then, the secondary antibody was added at 1:10,000 dilution in PBS-1% BSA buffer (100 µL per well) using either goat anti-mouse IgG-HRP, goat anti-mouse IgG1-HRP, goat anti-mouse IgG2a-HRP, or goat anti-mouse IgA-HRP (Thermo Fisher Scientific). After incubation for 1 h at 37°C and five washes with PBST buffer, plates were developed using the TMB (3,3’,5,5’-tetramethylbenzidine) Microwell Peroxidase Substrate System (KPL, 100 µL) for 5 to 10 min. The enzymatic reaction was stopped by adding 100 µL TMB BlueSTOP solution (KPL). The absorbance of optical density at 650 nm was read within 30 min on a VersaMax spectrophotometer. The endpoint titer was defined as the highest reciprocal dilution of serum that gives an absorbance more than twofold of the mean background of the assay.

### Virus neutralization assay

Neutralizing antibody titers in mouse immune sera against SARS-CoV-2 US-WA-1/2020 or its Beta, Delta, or Omicron variants were quantified by using Vero E6 cell–based microneutralization assay in the BSL-3 suite as previously described (Zhu et al., 2021). Briefly, serially 1:2 or 1:3 downward diluted mouse sera (original dilution 1:10 or 1:20) that were decomplemented at 56°C for 60 min in a 60 µl volume were incubated for 1 h at room temperature (RT) in duplicate wells of 96-well microtiter plates that contained 120 infectious SARS-CoV-2 virus particles in 60 µL in each well. After incubation, 100 µL of the mixture in individual wells was transferred to Vero E6 cell monolayer grown in 96-well microtiter plates containing 100 µL of MEM/2% fetal bovine serum (FBS) medium in each well and was cultured for 72 h at 37°C before assessing the presence or absence of cytopathic effect (CPE). Neutralizing antibody titers of the tested specimens were calculated as the reciprocal of the highest dilution of sera that completely inhibited virus-induced CPE.

### T cell proliferation and phenotypes, and cytokine analysis

To measure T-cell proliferation, bromodeoxyuridine (BrdU), a thymidine analog, incorporation method was used. Briefly, spleens were aseptically removed from 5 animals of each indicated group on day 21 after the last immunization dose. Spleens were homogenized and passed through a 70 µm cell strainer to obtain single cell suspension in RPMI 1640 cell culture medium. Splenocytes were then seeded into 24 well tissue culture plates at a density of 2.0 × 10^6^ cells/well (4 wells/mouse) and stimulated with either SARS-CoV-2 S-trimer (10-100 µg/mL) or SARS-CoV-2 PepTivator® Peptide S and NP protein Pools (10 µg/mL each, Miltenyi Biotec) for 72 h at 37°C. BrdU (BD Bioscience) was added to a final concentration of 10 µM during the last 18 h of incubation with the stimulants to be incorporated into the splenocytes (Endl et al., 1997; Penit, 1986). Subsequently, the BrdU-labeled splenocytes were surface stained for T-cell (CD3e-APC; eBioscience) marker after blocking with anti-mouse CD16/32 antibodies (BioLegend). Cells were then permeabilized and treated with DNase to expose BrdU epitopes followed by anti-BrdU-FITC and 7-AAD (7-amino-actinomycin D) staining by using BD Pharmingen FITC BrdU Flow Kit. The splenocytes were then subjected to flow cytometry, and data was analyzed as we previously described (Kilgore et al., 2021a; Kilgore et al., 2021b; Tiner et al., 2016). The percent of BrdU positive cells in CD3 positive populations were calculated using FACSDiva software.

To measure T-cell phenotypes, the above overnight (16 h) stimulated splenocytes were similarly blocked with anti-mouse CD16/32 antibodies (BioLegend) and stained with Fixable Viability Dye eFluor™ 506 (eBioscience) followed by APC anti-mouse CD3e (eBioscience), PE/Dazzle 594 anti-mouse CD4 (BioLegend), FITC anti-mouse CD8 (BioLegend) for CD3, CD4 and CD8 T-cell surface markers, respectively. Cells were then permeabilized for intracellular staining with PerCP/Cy5.5 anti-mouse IFNγ, PE/Cy7 anti-mouse IL-17A (BioLegend), eFluor 450 anti-mouse TNFα (eBioscience), and analyzed by flow cytometry.

To assess cytokine production, cell supernatants were collected after stimulation with S-trimers as described above for 72 h at 37°C. Cytokines in the supernatants were then measured by using Bio-Plex Pro mouse cytokine 23-plex assay (Bio-Rad Laboratories). Likewise BALF from control and immunized mice was used to measure cytokines.

### 16S rRNA gene sequencing and microbiome analysis

Fecal pellets were collected from 5 animals of each indicated group on day 21 after the last immunization dose. Total genomic DNA was extracted from the fecal matter using methods previously described (Salonen et al., 2010; Yu and Morrison, 2004). DNA samples were further purified using a DNA Clean and Concentrator kit (Zymo Research). The above extracted microbial DNA was then subjected to amplification and sequencing of the V4 region of the 16S rRNA gene by using a NEXTflex 16S V4 Amplicon Seq kit 2.0 (PerkinElmer), and sequences were generated on the Illumina MiSeq platform (Illumina). Raw reads were filtered using the Lotus pipeline (Hildebrand et al., 2014), followed by de novo clustering to operational taxonomic units (OTUs) at 97% sequence identity with UPARSE (Edgar, 2013). Bacterial diversity and community composition were evaluated using QIIME v1.8 (Caporaso et al., 2010), and taxonomy assignment of the representative sequence for each OTU was completed using the RDP classifier algorithm and the SILVA reference database (v123) (Quast et al., 2013).

### Animal challenges

Immunized and control mice were first ear tagged and their initial weights recorded. Mice were then anesthetized and intranasally challenged with 60 µl of either SARS-CoV-2 MA10 strain for conventional mice or SARS-CoV-2 US-WA-1/2020 strain or the Delta variant (B.1.617.2) for hACE2 transgenic mice. The challenge dose was ~10^5^ median tissue culture infectious dose (TCID_50_). For hACE2 transgenic mice, the challenge dose was 300 TCID_50_. The animals were monitored for the onset of morbidity (weight loss and other signs of illness, every day) and mortality over the indicated period.

### Histopathology Studies

Lung tissues were excised from euthanized animals (immunized and control) at 2-5 days post challenge and immersion fixed in 10% neutral buffered formalin. After fixation, tissues were sectioned at 5 µm, mounted on glass slides, and stained with hematoxylin and eosin (HE) and MOVAT for histopathological analysis (Department of Pathology, UTMB). Staining with MOVAT helps in better visualizing tissue architecture. Histopathological analysis of lung sections from Balb/c mice was performed based on three parameters: mononuclear inflammatory infiltrate around bronchovascular bundles, interstitial inflammation, and alveolar exudate/hemorrhage. Scores for bronchovascular infiltrates ranged from 0 (normal) to 3, as follow: 1-Occasional mononuclear infiltrates, 5-10 microns thick; 2: multifocal mononuclear infiltrates, 5-20 microns thick; and 3-Diffuse mononuclear infiltrates, > 20 microns thick. The scores for interstitial inflammation were as follow:1-occasional areas of widened alveolar septa; 2. Multifocal areas of widened alveolar septa; and 3-diffused widening of alveolar septa. For alveolar exudate/hemorrhage, the scores were: 1-occasional areas of alveolar exudate/hemorrhage; 2-multifocal areas of alveolar exudate/hemorrhage; and 3-diffused areas of alveolar exudate/hemorrhage. The combined scores for the vector control group and the T4-CoV-2 vaccine group were analyzed using the Student’s t-test.

For hACE2 transgenic mice, histopathological analysis was performed based on these parameters: interstitial inflammation/alveolar exudate and mononuclear inflammatory infiltrate around bronchovascular (BV) bundles. Interstitial inflammation/alveolar exudates were scored based on percentage of the lung surface area involved (0-100%), while scores for BV infiltrate ranged from 0 (normal) to 3 as follow: 1-occasional mononuclear infiltrates, 5-10 microns thick; 2-multifocal mononuclear infiltrates, 5-20 microns thick; and 3-Diffused mononuclear infiltrates, > 20 microns thick. The scores for the intranasal PBS control group, T4 vector control group, and the T4-CoV-2 vaccinated group were analyzed using the Student’s t-test if the groups passed the normality test (Shapiro-Wilk) or Mann-Whitney Rank sum test if the normality test failed.

### Statistics and software

Statistical analyses were performed by GraphPad Prism 9.0 software using one-way or two-way analysis of variance (ANOVA) with Tukey’s *post hoc* test or multiple t-test according to the generated data. We used Kaplan-Meier with log-rank (Mantel-Cox) test for animal survival studies. Significant differences between two groups were indicated by \**P*<0.05, \*\**P*<0.01, \*\*\**P*<0.001, and \*\*\*\**P*<0.0001. ns indicates not significant.

Photo credit: the mouse and immune cell images were created with BioRender.com. The figure data were organized by Photoshop CS6 (Adobe).

## ACKNOWLEDGMENTS

This research was supported by NIAID/NIH supplement grant 3R01AI095366-07S1 (subaward: 1100992-100) and in part by NIAID/NIH grants AI111538 and AI081726 and National Science Foundation grant MCB-0923873 to V.B.R. Special funding provided by the IHII-COVID19 pilot grant as well as the support through the John S. Dunn Endowed Chair to A.K.C. is greatly acknowledged.

## AUTHOR CONTRIBUTIONS

V.B.R. and A.K. C. designed and directed the project. J.Z. and V.B.R. designed vaccine constructs. S.J. designed Beta variant trimer and purification protocol. H.B. and S.J. purified WT and Secto-β trimers. N.A. produced vaccine phages. J.Z. prepared vaccine samples and performed all ELISAs and binding assays. P.B.K, V.T., E.K.H. performed animal studies; J.P.O. performed histopathology studies; A.K., C.L.G., and S.B. performed microbiota studies; A.D. and V.T. performed neutralization and viral load studies. J.Z., J.S., P.K.B., J.P.O., Y.M.H, A.K., C.L.G., S.B., A.D., V.T., C-Te. K.T., A.K.C., and V.B.R. analyzed and interpreted the data. V.B.R., A.K.C., and J.Z. wrote the manuscript.

## SUPPLEMENTAL INFORMATION

**Figure S1.**
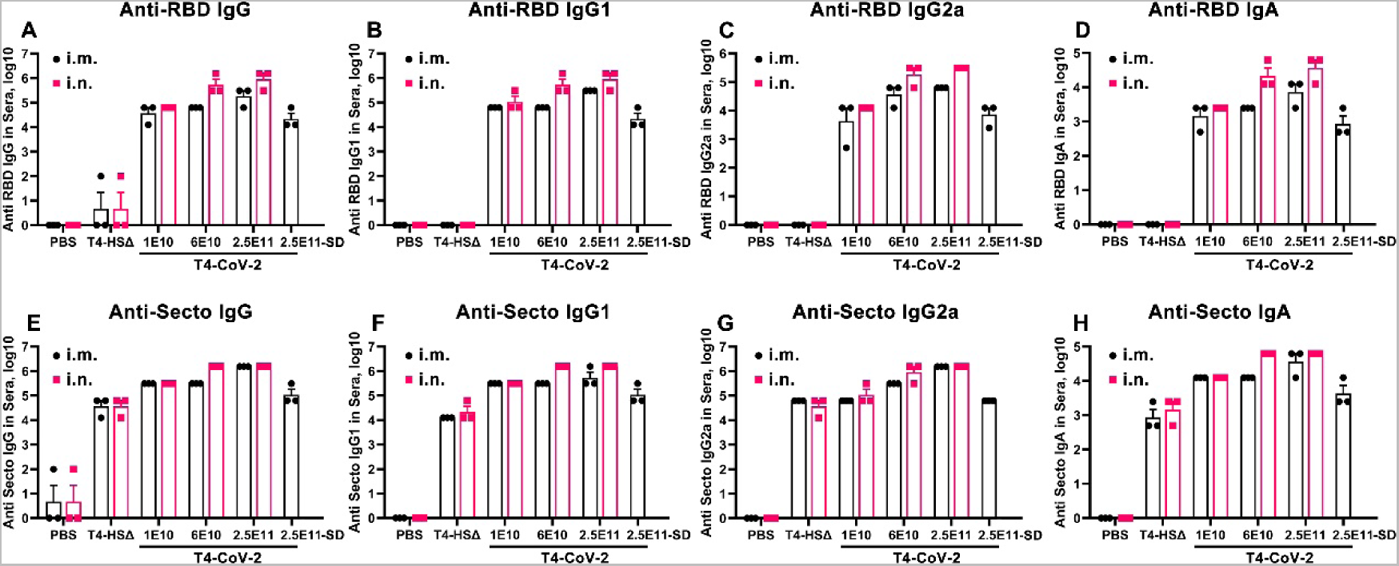
Anti-spike/RBD systemic humoral responses in sera from i.m. or i.n. administered BALB/c mice using various doses of T4-CoV-2 vaccine. ELISA assays were performed to measure reciprocal endpoint antibody titers of sera from i.m. (black) or i.n. (red) vaccinations: anti-RBD IgG **(A),** anti-RBD IgG1 **(B)**, anti-RBD IgG2a **(C)**, anti-RBD IgA **(D)**, anti-Secto IgG **(E)**, anti-Secto IgG1 **(F)**, anti-Secto IgG2a **(G)**, and anti-Secto IgA **(H)**. PBS and T4-HSΔ were used as naïve and vector controls, respectively. SD, single-dose. Data represent mean ± SEM. Data are from 3 pooled independent experiments (n = 22 for T4-CoV-2, n = 10 for T4-HSΔ, and n = 5 for PBS).

**Figure S2.**
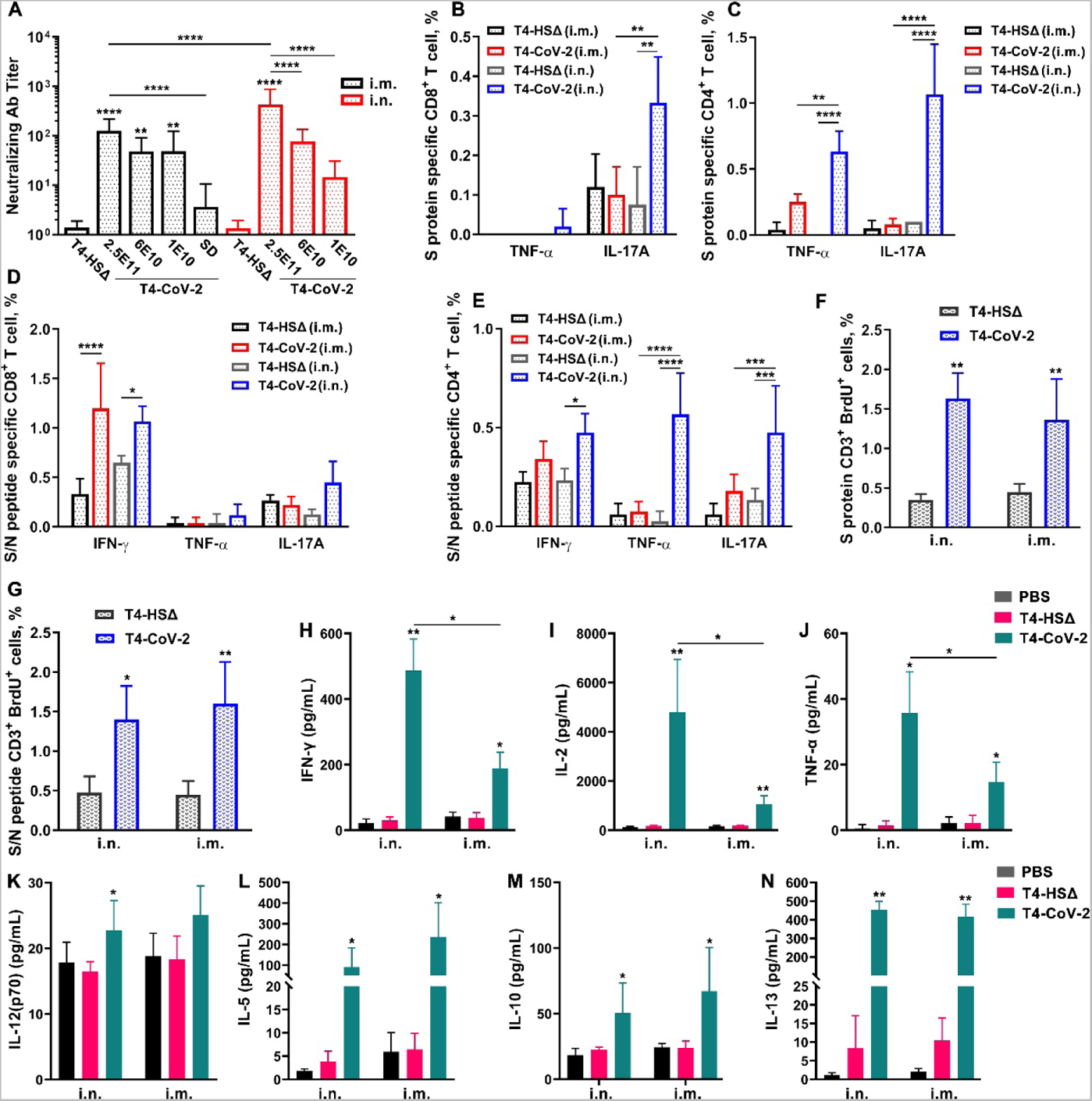
Neutralizing antibody and cellular immune responses in i.m. and i.n. vaccinated BALB/c mice. **(A)** Virus neutralizing activity in sera of i.m. and i.n. vaccinated mice was determined by Vero E6 cell cytopathic assay using ancestral SARS-CoV-2 US-WA-1/2020 strain. **(B and C)** Cellular immune responses after stimulation with purified Secto trimers. Cells were stained with T cell surface markers CD3, CD4, and CD8 followed by intracellular TNFα and IL-17A staining. Percentage of TNFα^+^ CD8^+^ (B), IL-17A^+^ CD8^+^ (B), TNFα^+^ CD4^+^ (C), IL-17A^+^ CD4^+^ (C) cells were plotted. **(D and E)** Cellular immune responses after stimulation with S- and NP-peptides. Percentage of IFNγ^+^ or TNFα^+^ or IL-17A^+^ in CD8^+^ (D) or CD4^+^ (E) cells were plotted. **(F and G)** T cell proliferation in mice immunized with T4-CoV-2 vaccine. Spleens were harvested from mice 21 days after the boost. Splenocytes were isolated and stimulated with either purified S protein trimer (F) or S- and NP-peptides (G). Cells were stained for T cell surface marker CD3 as well as for incorporated BrdU, and analyzed by flow cytometry. The percent of BrdU incorporation in CD3 positive cells was plotted. **(H to N)** Splenocyte cytokine response to S and NP peptide stimulation. The cytokines in the cell culture supernatants were analyzed by using Bioplex-23 assay. The representative Th1 (H to K) and Th2 (L to N) cytokines are shown. For A to G, data represent mean ± standard deviation and are representative of five biological replicates. A Two-way ANOVA with Tukey post hoc test was used; *P < 0.05; **P<0.01, ***P<0.001, ****P<0.0001. For H to N, statistical significance was determined using nonparametric Student’s t test compared to T4-HSΔ vector vs T4-CoV-2 groups, and i.n. vs i.m. administration groups; *P < 0.05; **P < 0.01. Data represent mean ± standard deviation and are representative of five biological replicates.

**Figure S3.**
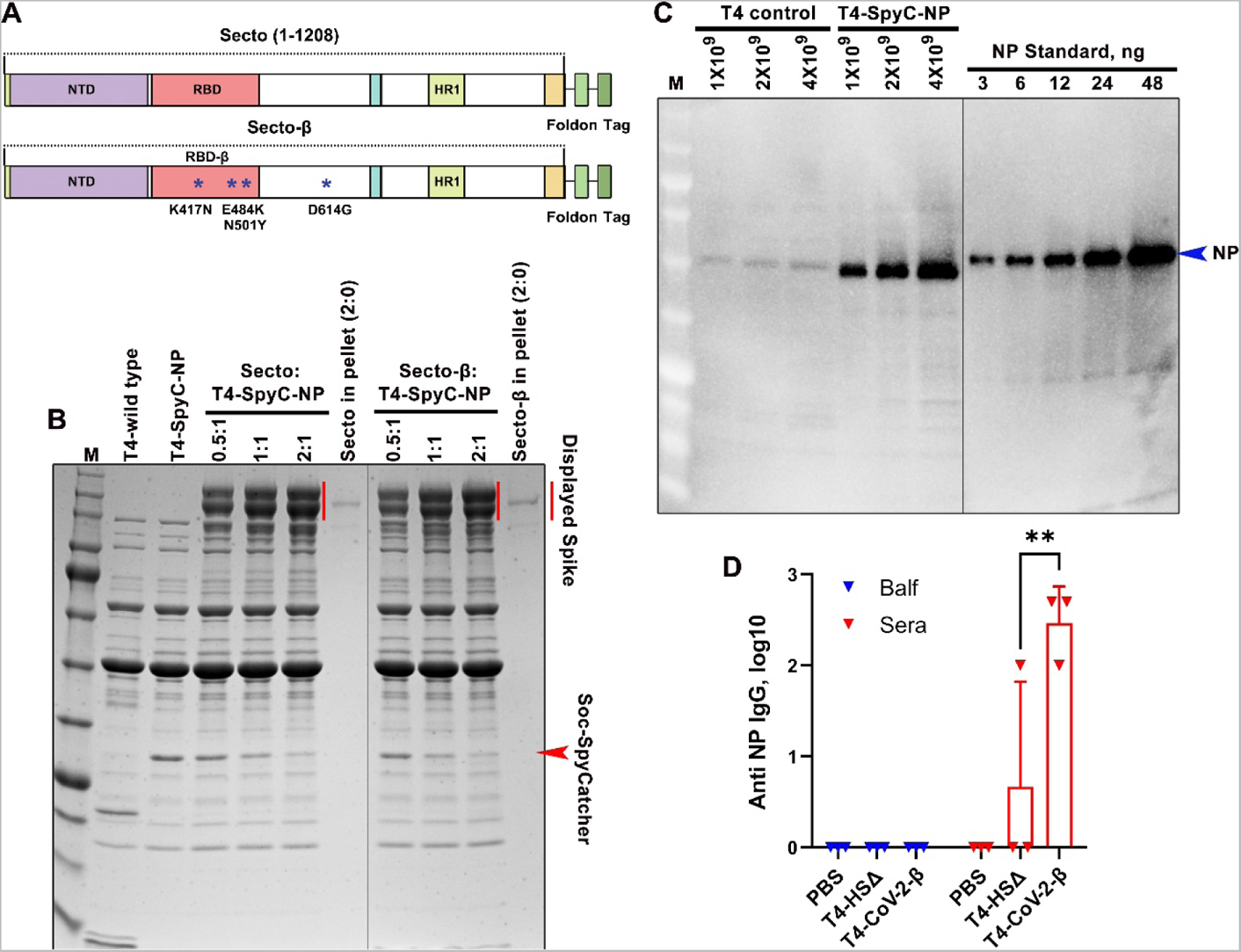
Characterization of Secto-β trimer, copy number of NP in T4-CoV-2-β vaccine, and anti-NP antibody responses. **(A)** Schematics of Secto and Secto-β cassette for expression in ExpiCHO cells. The Secto-β was constructed by incorporating K417N, E484K, and N501Y mutations in the receptor-binding domain (RBD) and D614G mutation in the S2 region of the trimer. These mutations are the core mutations of Beta variant which are responsible for the immune escape in vaccinated people. **(B)** *In vitro* display of Secto and Secto-β trimers on T4-SpyCatcher phage at increasing ratios of S protein molecules to Soc binding sites (0:1 to 2:1). S trimer and T4-SpyCatcher phage were incubated at 4°C for 1 h, followed by centrifugation to remove the unbound material. After two washes, the pellet was resuspended in buffer (which one!), and SDS-PAGE was performed. The positions of Soc-SpyCatcher (red arrowhead) and S protein bands (red line) are indicated. **(C)** Quantification of the copy number of NP protein molecules packaged in T4-CoV-2 vaccine by Western-blotting using commercial NP standard (Sino Biological). **(D)** Anti-NP antibody responses in sera (red) and BALF (blue) of immunized mice at day 21 after boosting. ELISA assays were performed to determine reciprocal endpoint antibody titers of anti-NP IgG. PBS and T4-HSΔ were used as naïve and vector controls, respectively. Data represent mean ± SEM. Data are from 3 pooled independent experiments (n = 15 for sera analysis and n = 5 for BALF analysis). P values were calculated using a Two-way ANOVA with Tukey post hoc test to compare multiple groups. **P<0.01.

**Figure S4.**
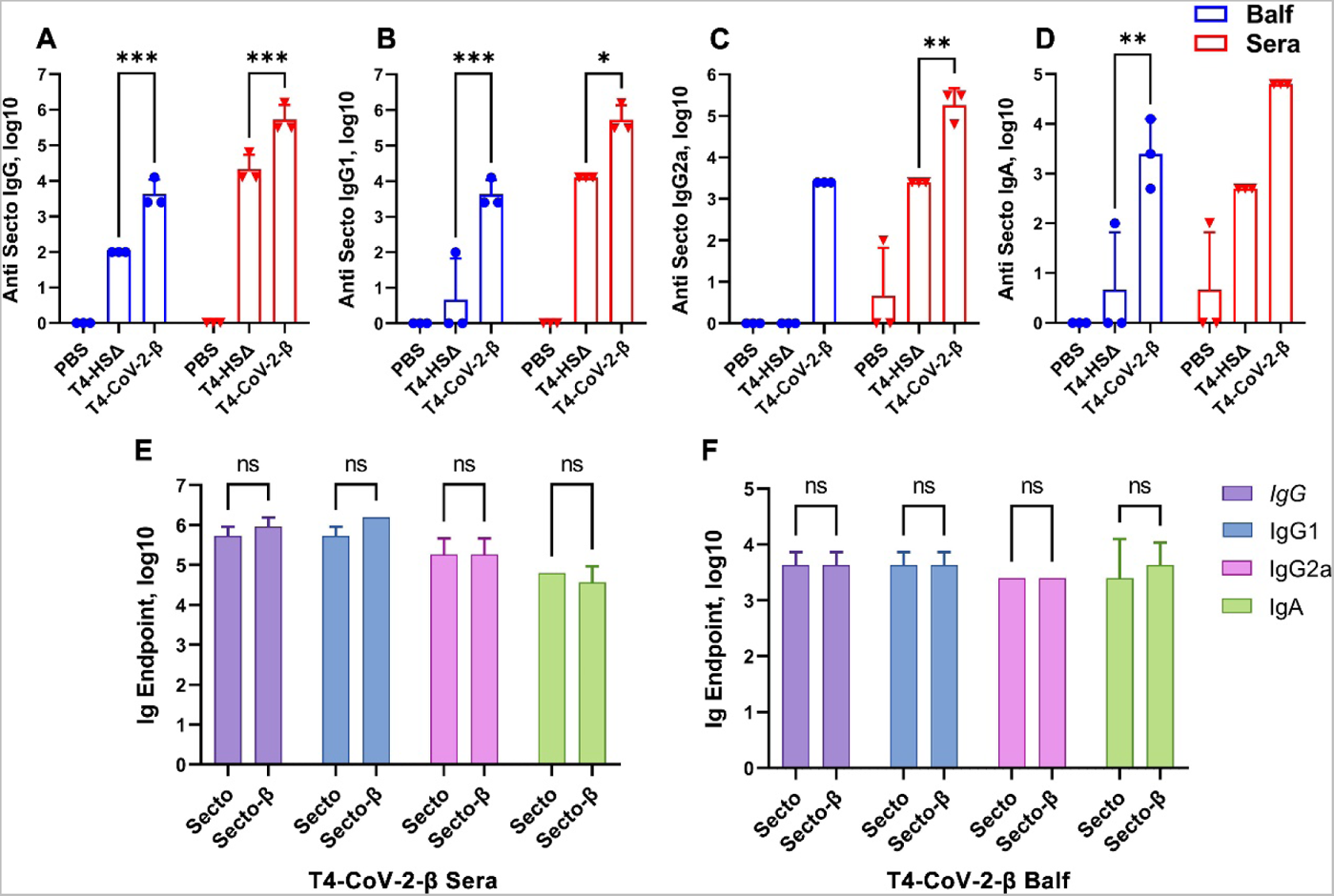
Secto- and Secto-β-trimer binding antibody titers in T4-CoV-2-β i.n. immunized hACE2 transgenic mice. **(A to D)** Antibody responses in sera (red) and BALF (blue) of immunized mice at day 21 after boosting. ELISA assays were performed to determine reciprocal endpoint antibody titers of anti-Secto IgG (A), anti-Secto IgG1 (B), anti-Secto IgG2a (C), and anti-Secto IgA (D). Data are from 3 pooled independent experiments (n = 15 for sera analysis and n = 5 for BALF analysis). P values were calculated using a Two-way ANOVA with Tukey post hoc test to compare multiple groups. *P<0.05, **P<0.01, ***P<0.001. **(E and F)** Comparison between anti-Secto and anti-Secto-β antibody responses (IgG, IgG1, IgG2a, and IgA) in sera (E) and BALF (F). ns, no significance.

**Figure S5.**
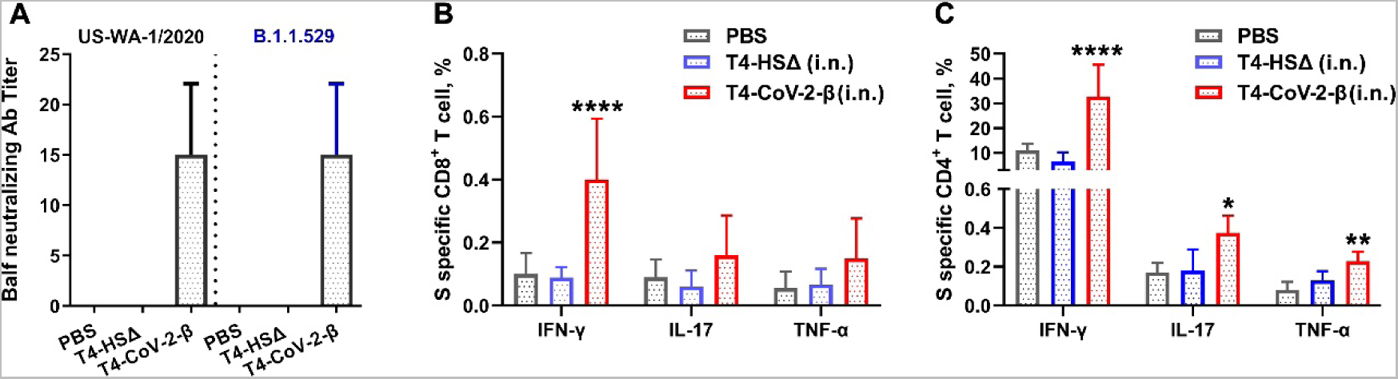
Virus neutralization activity in BALF and T cell immune responses in hACE2 transgenic mice. **(A)** Neutralizing antibody titers in BALF were determined by Vero E6 cell cytopathic assay using ancestral SARS-CoV-2 US-WA-1/2020 and B.1.1.529 (Omicron) strains. Data represent mean ± standard deviation. Data are from 2 pooled replicate experiments. **(B and C)** Analysis of CD8^+^ (B) and CD4^+^ (C) T cell immune response in stimulation with Secto-β protein. Cells were then stained with T cell surface markers CD3, CD4, and CD8 followed by intracellular IFNγ, TNFα and IL-17A staining. Percentage of IFNγ^+^ or TNFα^+^ or IL-17A^+^ in CD4^+^ or CD8^+^ cells were plotted. P values were calculated using a Two-way ANOVA with Tukey post hoc test to compare multiple groups; *P < 0.05; **P<0.01, ****P<0.0001. Data represent mean ± standard deviation. Data are representative of five biological replicates.

